# CRISPR activation rescues abnormalities in *SCN2A* haploinsufficiency-associated autism spectrum disorder

**DOI:** 10.1101/2022.03.30.486483

**Authors:** Serena Tamura, Andrew D. Nelson, Perry W.E. Spratt, Henry Kyoung, Xujia Zhou, Zizheng Li, Jingjing Zhao, Stephanie S. Holden, Atehsa Sahagun, Caroline M. Keeshen, Congyi Lu, Elizabeth C. Hamada, Roy Ben-Shalom, Jen Q. Pan, Jeanne T. Paz, Stephan J. Sanders, Navneet Matharu, Nadav Ahituv, Kevin J. Bender

## Abstract

The majority of autism spectrum disorder (ASD) risk genes are associated with ASD due to haploinsufficiency, where only one gene copy is functional. Here, using *SCN2A* haploinsufficiency, a major risk factor for ASD, we show that increasing the expression of the existing functional *SCN2A* allele with CRISPR activation (CRISPRa) can provide a viable therapeutic approach. We first demonstrate therapeutic potential by showing that restoring *Scn2a* expression in adolescent heterozygous *Scn2a* conditional knock-in mice rescues electrophysiological deficits associated with *Scn2a* haploinsufficiency. Next, using an rAAV-CRISPRa based treatment, we restore electrophysiological deficits in both *Scn2a* heterozygous mice and human stem-cell-derived neurons. Our results provide a novel therapeutic approach for numerous ASD-associated genes and demonstrate that rescue of *Scn2a* haploinsufficiency, even at adolescent stages, can ameliorate neurodevelopmental phenotypes.

## INTRODUCTION

Autism spectrum disorder (ASD) is a common neurodevelopmental disorder (NDD) affecting 1 in 44 children eight years of age or approximately 2.5% of the general population in the US (Maenner, 2021). Recent advances in gene discovery have facilitated the identification of 255 high confidence ASD *de novo* risk genes (Fu et al., 2021). One hundred sixty-one of these risk genes are either known or predicted to cause ASD due to haploinsufficiency, where the mRNA/protein levels from the residual gene copy are insufficient to enable the typical function of the gene (Karczewski et al., 2020). Delivering functional gene copies via gene therapy could potentially restore transcriptional balance and rectify deficits in haploinsufficient diseases. Gene therapy relies primarily on using recombinant adeno-associated virus (rAAV) for transgene delivery due to its limited pathogenicity and long-term transgene expression (Wang et al., 2019). However, rAAV has a limited packaging capacity (4,700 base pairs optimal packaging capacity minus ∼1,700 base pairs needed for transgene expression) (Wu et al., 2010). Examination of the coding sequence (CDS) length of the 161 likely haploinsufficient ASD risk genes shows that 77 exceed rAAV vector capacity (**Table S1**), excluding them from traditional gene replacement therapy approaches (Howe et al., 2021).

Here, we tested whether *cis*-regulation therapy (CRT) could be a viable approach for treating ASD-associated haploinsufficiency. CRT utilizes a nuclease-deficient gene-editing system, such as a “dead” Cas9 (dCas9), fused to a transcriptional modulator to target gene regulatory elements (i.e. promoter or enhancer) and alter gene expression (Matharu and Ahituv, 2020). For this study, we used dCas9 fused to a transcriptional activator, termed CRISPR activation (CRISPRa), to upregulate gene expression and restore haploinsufficiency by targeting a gene promoter. For our proof-of-concept, we utilized ASD-associated haploinsufficiency of the sodium voltage-gated channel alpha subunit 2 (*SCN2A*), a gene with a six kilobase-long CDS (Howe et al., 2021). Variants in this gene can enhance channel function, resulting in seizure phenotypes, or inhibit channel function, resulting in ASD (Sanders et al., 2018). Loss-of-function (LoF) variants in *SCN2A* are one of the most significant genetically-defined subsets of ASD, second to Fragile X syndrome (Ben-Shalom et al., 2017), and contribute to over 10,000 cases of ASD in the US per year (0.3% of all ASD cases) (Satterstrom et al., 2020). In addition to ASD, individuals frequently have severe intellectual disability and poor developmental outcomes (Wolff et al., 2017).

*SCN2A* encodes the neuronal voltage-gated sodium channel, Na_V_1.2, which is broadly expressed in the central nervous system, including in neocortical excitatory pyramidal cells whose dysfunction is implicated in ASD etiology (Willsey et al., 2013). Within the first year of life in humans and before postnatal (P) day 7 in mice, Na_V_1.2 plays an essential role as the primary Na_V_ responsible for action potential (AP) initiation in pyramidal cell axons (Gazina et al., 2015). After this period, Na_V_1.2 is enriched throughout the somatic and dendritic domains in addition to the most proximal region of the axon initial segment (Hu et al., 2009; Spratt et al., 2019). *Scn2a* haploinsufficiency in mice—either from birth (constitutive) or conditionally induced after P7—results in impairments in dendritic intrinsic excitability and excitatory synapse function (**Fig. 1A**) (Spratt et al., 2021, 2019). This suggests that Na_V_1.2 actively helps maintain dendritic function throughout life and raises the possibility that rescue from *Scn2a* haploinsufficiency can be therapeutic, even if such a rescue is administered later in life.

**Figure 1.**
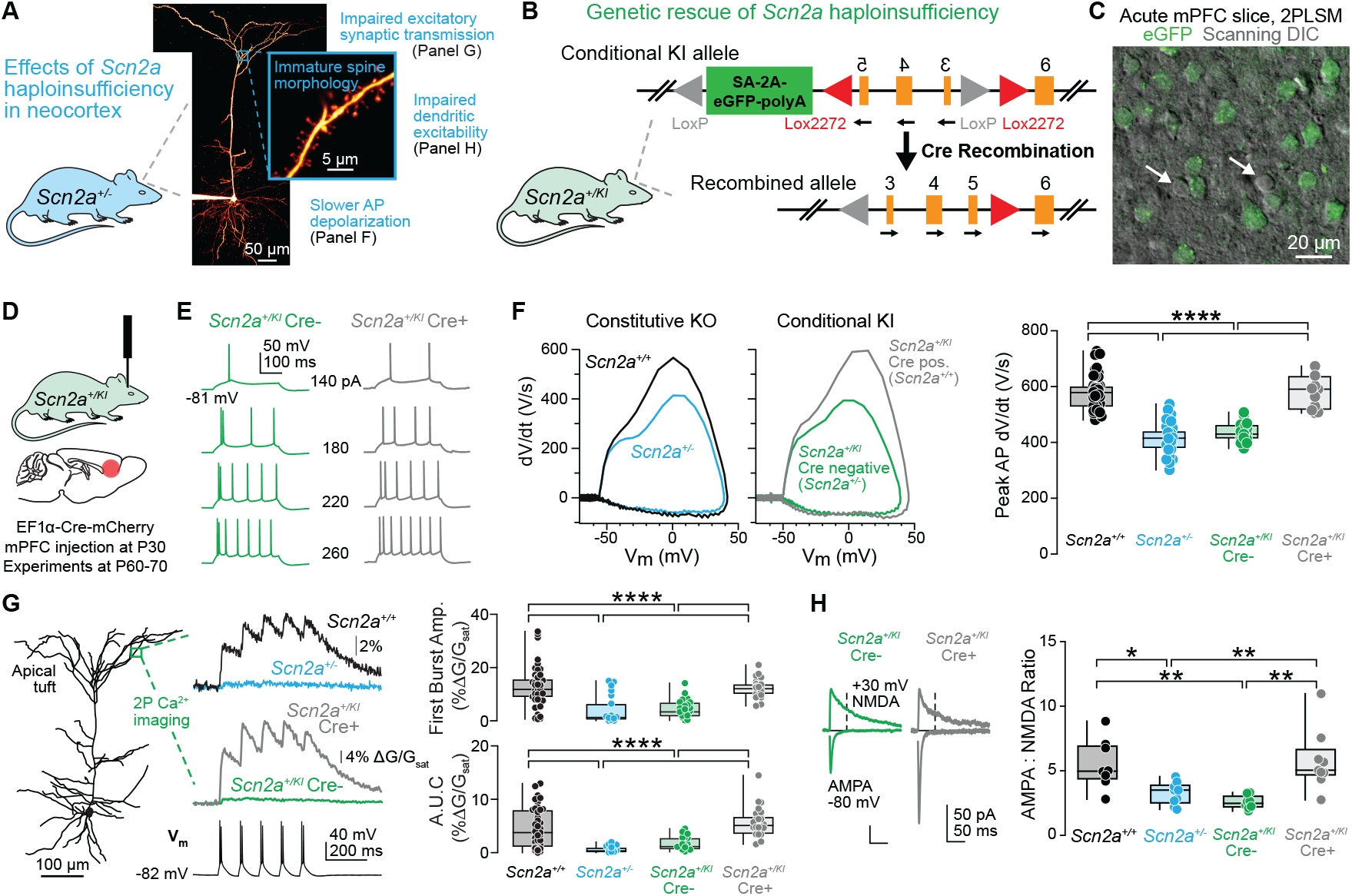
Genetic rescue from *Scn2a* haploinsufficiency in adolescent mice rescues electrophysiological deficits. **A:** Summary of *Scn2a* haploinsufficiency effects on mouse neocortical layer 5 thick-tufted neuronal physiology and anatomy. **B:** Genetic design and strategy of genetic restoration of *Scn2a* in the *Scn2a*^*+/KI*^ conditional knock-in mouse model. **C:** 2PLSM z-stack of layer 5b medial prefrontal cortex (mPFC) from a coronal brain slice from *Scn2a*^*+/KI*^ inducible recovery mouse without Cre injection. Arrows highlight GFP-negative, presumptive interneurons. **D:** Schematic of rAAV-EF1α-Cre-mCherry injection into the mPFC of P30 *Scn2a*^*+/KI*^ mice. Electrophysiological and imaging experiments in *Scn2a*^*+/KI*^ and control mice were performed between P60-70. **E:** APs per 300 ms stimulation epoch across a range of current amplitudes in *Scn2a*^*+/KI*^ Cre-negative (green) and *Scn2a*^*+/KI*^ Cre positive (gray) layer 5 pyramidal neurons. **F:** Left: Representative phase-plane plots (dV/dt vs. voltage) of somatic APs in *Scn2a*^*+/+*^ (black), *Scn2a*^*+/-*^ (cyan), *Scn2a*^*+/KI*^ Cre-(green) and *Scn2a*^*+/KI*^ Cre+ (gray) neurons. Right: Peak dV/dt of the first AP evoked by a near-rheobase current. Circles represent single cells. *Scn2a*^*+/+*^: 578.2 ± 9.3 V/s, n = 44; *Scn2a*^*+/-*^: 413.9 8.8, n = 37, *Scn2a*^*+/KI*^ Cre-: 435.4 ± 8.1 n = 19; *Scn2a*^*+/KI*^ Cre+: 578.4 ± 17.7, n = 11. *Scn2a*^*+/+*^ vs. *Scn2a*^*+/-*^: ****p < 0.0001, *Scn2a*^*+/+*^ vs. *Scn2a*^*+/KI*^ Cre-: ****p < 0.0001, *Scn2a*^*+/-*^ vs. *Scn2a*^*+/KI*^ Cre+: ****p < 0.0001, *Scn2a*^*+/KI*^ Cre-vs. *Scn2a*^*+/KI*^ Cre+: ****p < 0.0001. Holm-Šídák multiple comparisons test. **G:** Left: 2PLSM imaging of bAP-evoked Ca transients in the apical dendrites (>400 μm from soma) of layer 5b neurons in P60-70 *Scn2a*^*+/+*^, *Scn2a*^*+/-*^, *Scn2a*^*+/KI*^ Cre-, *Scn2a*^*+/KI*^ Cre+ mice. Calcium transients were evoked by bursts of AP doublets. Right: Ca transient amplitude of the first of 5 bursts (top) and area under the curve (AUC) (bottom). Circles represent imaging sites. First burst amplitude: *Scn2a*^*+/+*^: 12.7 ± 1.0, n = 49 sites; *Scn2a*^*+/-*^: 3.7 ± 1.4, n = 23 sites, *Scn2a*^*+/KI*^ Cre-: 4.4 ± 0.7 n = 29 sites; *Scn2a*^*+/KI*^ Cre+: 12.2 ± 0.7, n = 26 sites. AUC: *Scn2a*^*+/+*^: 4.5 ± 0.5, n = 49 sites; *Scn2a*^*+/-*^: 0.5 ± 0.1, n = 23 sites, *Scn2a*^*+/KI*^ Cre-: 1.6 ± 0.2 n = 29 sites; *Scn2a*^*+/KI*^ Cre+: 5.7 ± 0.5, n = 26 sites. First burst amplitude and AUC: *Scn2a*^*+/+*^ vs. *Scn2a*^*+/-*^: ****p < 0.0001, *Scn2a*^*+/+*^ vs. *Scn2a*^*+/KI*^ Cre-: ****p < 0.0001, *Scn2a*^*+/-*^ vs. *Scn2a*^*+/KI*^ Cre+: ****p < 0.0001, *Scn2a*^*+/KI*^ Cre-vs. *Scn2a*^*+/KI*^ Cre+: ****p < 0.0001. Holm-Šídák multiple comparisons test. **H:** Left: AMPA receptor-mediated and mixed AMPA/NMDA receptor-mediated evoked EPSCs at −80 and +30 mV, respectively. NMDA receptor-mediated component was calculated 50 ms after stimulation (dotted line). Right: Quantification of AMPA:NMDA ratio. *Scn2a*^*+/+*^: 5.5 ± 0.7, n = 8 cells; *Scn2a*^*+/-*^: 3.3 0.3, n = 9 cells, *Scn2a*^*+/KI*^ Cre-: 2.6 ± 0.2 n = 8 cells; *Scn2a*^*+/KI*^ Cre+: 5.8 ± 0.8, n = 9 cells. *Scn2a*^*+/+*^ vs. *Scn2a*^*+/-*^: *p = 0.02, *Scn2a*^*+/+*^ vs. *Scn2a*^*+/KI*^ Cre-: **p = 0.004, *Scn2a*^*+/-*^ vs. *Scn2a*^*+/KI*^ Cre+: **p = 0.009, *Scn2a*^*+/ KI*^ Cre-vs. *Scn2a*^*+/KI*^ Cre+: **p = 0.002. Holm-Šídák multiple comparisons test.

As the intrinsic and synaptic deficits of *SCN2A* haploinsufficiency can be readily measured via electrophysiology, we hypothesized that *SCN2A* could provide an efficient and quantifiable model for CRT-based interventions. We administered rAAV-based CRISPRa targeting the *Scn2a* promoter to *Scn2a* heterozygous mice and neurons differentiated from human embryonic stem cells (hESCs) (Lu et al., 2019) and demonstrate that this approach can rescue excitability deficits in both models. In mice, we also observed that CRISPRa can rescue these deficits at adolescent stages, and does not increase seizure risk. Combined, our work showcases a potential therapeutic approach that could be applied to many genes that when haploinsufficient are associated with ASD risk and suggests that rescue later in life could ameliorate electrophysiological phenotypes associated with *Scn2a* haploinsufficiency.

## RESULTS

### Restoration of two functional *Scn2a* copies in adolescent mice rescues electrophysiological deficits

To test the feasibility of increasing functional gene copy number to rescue the electrophysiological deficits associated with *Scn2a* haploinsufficiency, we first developed a conditional knock-in mouse line. In this mouse, termed *Scn2a*^*+/KI*^, exons 3-5 of one *Scn2a* allele were flipped and flanked by LoxP sites, with an eGFP sequence in frame to visualize cells that express *Scn2a* (**Fig. 1B**). GFP-positive neurons targeted for whole recordings had electrophysiological features indicative of *Scn2a*^*+/-*^ pyramidal cells, including a decrease in the peak change in voltage during AP depolarization (peak dV/dt) relative to *Scn2a*^*+/+*^ controls (**Fig. 1C-F**). Moreover, parvalbumin-positive interneurons were GFP negative, consistent with known Na_V_1.2 functional expression patterns (**Fig. S1**) (Li et al., 2014; Spratt et al., 2019).

A previous study showed that inducing *Scn2a* haploinsufficiency during adolescence impairs intrinsic and synaptic function (Spratt et al., 2019). To test whether these deficits can be rescued by gene reinstatement during adolescence, we injected rAAV-EF1α-Cre-mCherry into the medial prefrontal cortex (mPFC) of these *Scn2a*^*+/KI*^ mice around P30 (**Fig. 1D**). Four weeks post injection, mCherry-positive neurons were targeted for whole-cell recordings in acute slices. In the presence of Cre, multiple measures of neuronal and synaptic function were comparable to those found in wild-type (WT) mice, including peak AP dV/dt, AP-burst evoked calcium influx in the apical dendritic tuft, and AMPA:NMDA ratio (**Fig. 1F-H**). Together, these data show that features of intrinsic and synaptic function that depend directly and indirectly on proper Na_V_1.2 function can be restored to WT levels upon reactivation of the second allele in adolescent stages. They also suggest that the development of a CRT approach to increase expression of the functional *SCN2A* allele could rescue these deficits caused by haploinsufficiency.

### *Scn2a-CRISPRa* optimization *in vitro*

To optimize an rAAV-based CRISPRa approach to rescue *Scn2a* haploinsufficiency, nine different sgRNAs targeting the *Scn2a* promoter were screened for their ability to upregulate *Scn2a* in mouse neuroblastoma cells (Neuro-2a). Guides were co-transfected with a *Staphylococcus aureus* (sa) dCas9 fused to the transcriptional activator VP64 (**Table S2**) (Flint and Shenk, 1997). sadCas9-VP64 is compact in size (3.3 kb), allowing it to fit into an rAAV vector, and has been shown to promote upregulation to near WT levels in other forms of haploinsufficiency (Matharu et al., 2019). Three guides increased *Scn2a* mRNA levels and were packaged into rAAV-DJ serotype, which provides high expression levels in many tissues (Grimm et al., 2008). rAAV-sgRNA viruses were then co-transduced with the rAAV-sadCas9-VP64 activator into Neuro-2a cells for five days. One of the three rAAV-sgRNA was found to significantly increase *Scn2a* mRNA expression by two-fold and was selected for subsequent mouse studies (**Fig. S2A-B**). To test for off-target effects, we used qPCR and RNA-seq to measure the mRNA expression of all sodium channels within the same topologically associated domain (TAD) following plasmid CRISPRa transfection into Neuro-2a cells. We observed that *Scn2a* was the only significantly upregulated central nervous system (CNS) related sodium channel compared to cells transfected with a no sgRNA negative control (**Fig. S2C-D**).

### Administration of CRISPRa restores cell-autonomous excitability in *Scn2a*+/- mice

To test whether *Scn2a*-rAAV-CRISPRa (rAAV-sadCas9-VP64 + rAAV-*Scn2a*-sgRNA) could be used as a therapeutic intervention for *Scn2a* haploinsufficiency in adolescent mice, we stereotaxically injected *Scn2a*-rAAV-CRISPRa into the mPFC of constitutive *Scn2a*^+/-^ mice at approximately P30. Mice were injected unilaterally in one hemisphere and the opposite hemisphere was used as a biologically-matched uninjected control (**Fig. 2A-B**). As a negative (no sgRNA) control, *Scn2a*^+/-^ mice were co-injected with rAAV-sadCas9-VP64 along with an rAAV-mCherry (termed Empty Vector). Four weeks post-injection, the mPFC of the injected and uninjected hemispheres were dissected and extracted for qPCR analysis and electrophysiological recordings. *Scn2a*^+/-^ mice injected with *Scn2a*-rAAV-CRISPRa showed an upregulation of *Scn2a* mRNA of around 1.5-fold compared to the uninjected hemisphere, while rAAV-empty-mCherry injected mice showed no upregulation of *Scn2a* mRNA expression (**Fig. 2C**).

**Figure 2.**
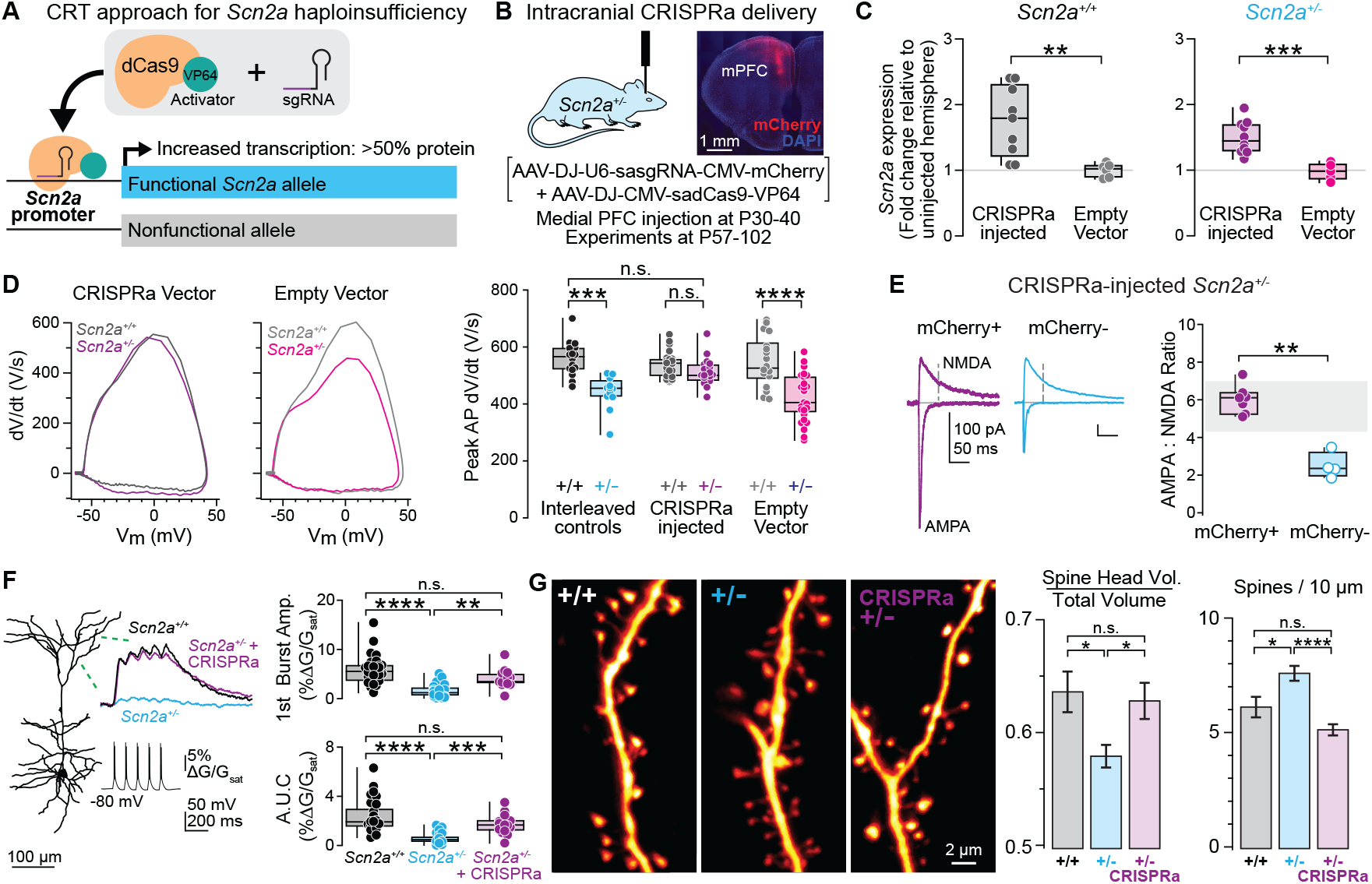
Intracranial *Scn2a*-rAAV-CRISPRa in mPFC rescues excitability deficits in *Scn2a*^+/-^ neurons. **A:** Design for *cis*-regulatory therapy using CRISPRa to rescue *Scn2a* haploinsufficiency. **B:** Left: Cartoon of local injection of *Scn2a*-rAAV-CRISPRa, which consists of both AAV-DJ-U6-sasgRNA-CMV-mCherry and AAV-DJ-CMV-sadCas9-VP64 (in brackets) into the mPFC of P30-P40 *Scn2a*^+/+^ or *Scn2a*^+/-^ mice. Right: Confocal image of coronal brain section immunostained with anti-mCherry and DAPI following unilateral injection of *Scn2a*-rAAV-CRISPRa in mPFC. **C:** qPCR of *Scn2a* mRNA of hemisphere injected with *Scn2a*-rAAV-CRISPRa or *Scn2a-*rAAV-empty normalized to uninjected hemisphere of P57-85 *Scn2a*^+/+^ (left) or *Scn2a*^+/-^ (right) mice. Circles represent single mice. *Scn2a*^+/+^ + CRISPRa: 1.7 ± 0.2, n = 9 mice; *Scn2a*^+/+^ + empty: 1.0 ± 0.04, n = 6 mice. **p = 0.0016. Mann-Whitney test. *Scn2a*^+/-^ + CRISPRa: 1.5 ± 0.08, n = 10 mice; *Scn2a*^+/-^ + empty: 1.0 ± 0.06, n = 5 mice. ***p = 0.0007. Mann-Whitney test. **D:** Left: Representative phase-plane plots of somatic APs from P57-85 *Scn2a*^+/+^ and *Scn2a*^*+/-*^ mice injected with *Scn2a*-rAAV-CRISPRa or *Scn2a-*rAAV-empty in mPFC. Right: Quantification of peak AP dV/dt. Circles represent single cells. *Scn2a*^+/+^: 559.8 ± 14.9, n = 16 cells; *Scn2a*^+/-^: 441.5 ± 17.1 n = 12 cells; *Scn2a*^+/+^ + CRISPRa: 539.2 ± 9.1, n = 24; *Scn2a*^+/-^ + CRISPRa: 509.1 ± 11.8, n = 19; *Scn2a*^+/+^ + empty: 548.5 ± 20.7, n = 18; *Scn2a*^+/-^ + empty: 432.7 ± 14.9, n = 29. *Scn2a*^+/+^ vs. *Scn2a*^+/-^ + empty: ****p < 0.0001, *Scn2a*^+/-^ vs. *Scn2a*^+/+^ + CRISPRa: ***p = 0.0006, *Scn2a*^+/-^ vs. *Scn2a*^+/+^ + empty: ***p = 0.0003, *Scn2a*^+/+^ + CRISPRa vs. *Scn2a*^+/-^ + empty: ****p < 0.0001. *Scn2a*^+/-^ + CRISPRa vs. *Scn2a*^+/-^ + empty: ***p = 0.0003, *Scn2a*^+/+^ + empty vs. *Scn2a*^+/-^ + empty: ****p < 0.0001. Holm-Šídák multiple comparisons test. **E:** Left: AMPA receptor-mediated and mixed AMPA/NMDA receptor-mediated evoked EPSCs at −80 and +30 mV, respectively in *Scn2a*^+/-^ + CRISPRa (mCherry-positive) (purple) neurons compared to *Scn2a*^+/-^ (mCherry-negative) (cyan) internal control cells. NMDA receptor-mediated component was calculated 50 ms after stimulation (dotted line). Right: Quantification of AMPA:NMDA ratio. Circles represent single cells. mCherry+: 6.0 0.3, n = 7; mCherry-: 2.5 ± 0.3, n = 4. Gray bar represents 95% confidence interval of *Scn2a*^+/+^ control data from Spratt 2019. mCherry+ vs. mCherry-: **p = 0.006. Mann-Whitney Wilcoxon test. **F:** Left: 2PLSM calcium imaging of bAP-evoked Ca transients in the apical dendrites (>400 um from soma) of layer 5b neurons from P55-64 *Scn2a*^+/+^ (black), *Scn2a*^+/-^ (cyan), and *Scn2a*^+/-^ + CRISPRa (purple) mice. Calcium transients were evoked by bursts of AP doublets. Right: Ca transient amplitude of the first of 5 bursts (top) and area under the curve (AUC) (bottom). Circles represent imaging sites from 3-8 cells per group. First burst amplitude: *Scn2a*^*+/+*^: 5.9 ± 0.6, n = 29 sites; *Scn2a*^*+/-*^: 1.6 ± 0.3, n = 28 sites, *Scn2a*^+/-^ + CRISPRa: 4.1 ± 0.5 n = 14 sites. *Scn2a*^*+/+*^ vs. *Scn2a*^*+/-*^ ****p < 0.0001, *Scn2a*^*+/-*^ vs. *Scn2a*^+/-^ + CRISPRa **p = 0.0018. Kruskal-Wallis test. AUC: *Scn2a*^*+/+*^: 2.3 ± 0.3, n = 29 sites; *Scn2a*^*+/-*^: 0.6 ± 0.1, n = 28 sites, *Scn2a*^+/-^ + CRISPRa: 1.7 ± 0.2 n = 14 sites. *Scn2a*^*+/+*^ vs. *Scn2a*^*+/-*^ ****p < 0.0001, *Scn2a*^*+/-*^ vs. *Scn2a*^+/-^ + CRISPRa ***p = 0.0002. Kruskal-Wallis test. **G:** Left: 2PLSM z-stacks of dendritic spines on apical dendritic tuft shafts of layer 5b neurons from P88-102 *Scn2a*^+/+^ (black), *Scn2a*^+/-^ (cyan), and *Scn2a*^+/-^ + CRISPRa (purple) mice. Right: Quantification of volume of the spine head relative to total volume of the head and shaft and number of spines per length (10 μm) of dendrite (mean ± SEM). Spine head volume: *Scn2a*^+/+^: 0.64 ± 0.02 (n = 226 spines, 13 branches, 2 mice); *Scn2a*^+/-^: 0.58 0.01 (n = 619 spines, 26 branches, 2 mice); *Scn2a*^+/-^ + CRISPRa: 0.63 ± 0.02 (n = 261 spines, 10 branches, 2 mice). *Scn2a*^+/+^ vs. *Scn2a*^+/-^: *p = 0.02, *Scn2a*^+/-^ vs. *Scn2a*^+/-^ + CRISPRa: *p = 0.03. Holm-Šídák multiple comparisons test. Spine density: *Scn2a*^+/+^: 6.1 ± 0.5 spines/10 m; *Scn2a*^+/-^: 7.6 ± 0.3 spines/10 m; *Scn2a*^+/-^ + CRISPRa: 5.1 ± 0.2 spines/10 m. *Scn2a*^+/+^ vs. *Scn2a*^+/-^: *p = 0.02, *Scn2a*^+/-^ vs. *Scn2a*^+/-^ + CRISPRa: ****p < 0.0001. Holm-Šídák multiple comparisons test.

To assess the phenotypic effects of *Scn2a*-rAAV-CRISPRa on neuronal excitability, acute slices from the mPFC were dissected four weeks post-injection and electrophysiological recordings were carried out from neurons in these slices. AP waveform, measured with somatic current-clamp recordings, are sensitive to *Scn2a* expression levels in single cells, with peak dV/dt being directly proportional to Na_V_1.2 density at the somatic membrane (Spratt et al., 2021, 2019). We observed that peak dV/dt was restored to WT levels in mCherry-positive neurons in CRISPRa-treated *Scn2a*^*+/-*^ mice (**Fig. 2D**). mCherry-negative neurons, recorded either in the same slices or in interleaved constitutive controls, displayed dV/dt values expected for *Scn2a*^*+/+*^ and *Scn2a*^*+/-*^ neurons respectively (**Fig. 2D**). Similarly, no upregulation in dV/dt was noted in mCherry-positive empty vector controls (**Fig. 2D**).

*Scn2a* haploinsufficiency is also known to impair dendritic excitability and excitatory synapse function. We found that *Scn2a*-rAAV-CRISPRa administration around P30 was able to rescue these impairments. AMPA:NMDA ratio, which is a common proxy for synapse strength and is abnormally low in *Scn2a*^*+/-*^ cells (Spratt et al., 2019), was rescued to WT levels (**Fig. 2E**). Similarly, AP-evoked calcium transients in distal apical dendrites, which are readily observed in WT but not *Scn2a*^*+/-*^ cells, were also restored (**Fig. 2F**). Furthermore, spine morphology alterations, including relative spine head size and density, were also restored to WT levels (**Fig. 2G**). Of note, the increase in overall spine density in *Scn2a*^*+/-*^ cells observed here, at P88-102, was not observed in previous work that focused on younger mice (P24-36) (Spratt et al., 2019). This suggests that mPFC *Scn2a*^*+/-*^ cells do not experience as much spine pruning and refinement as would normally occur during adolescence and early adulthood (Caballero et al., 2021), and that CRT for *Scn2a* may help re-engage such processes.

### Systemic CRT rescues *Scn2a+/-* electrophysiological deficits

We next explored the feasibility of intravenous systemic rAAV-CRISPRa delivery, as it is a more clinically translatable delivery approach. *Scn2a*^+/-^ mice were tail vein injected with *Scn2a*-rAAV-CRISPRa constructs packaged using the PhP.eb rAAV serotype, which can readily pass through the blood brain barrier in C57BL/6 mice and provide robust brain transgene expression (Deverman et al., 2016). Mice were injected at P30 and after four weeks showed widespread expression throughout the brain (**Fig. 3A-B**). qPCR showed high expression levels of *dCas9* and *mCherry* mRNA expression only in neocortical samples from tail-vein-treated *Scn2a*^*+/-*^ mice (**Fig. S2E**). Electrophysiological analysis of *Scn2a*-rAAV-CRISPRa-PhP. eb tail vein injected *Scn2a*^+/-^ mice showed a rescue of AP peak dV/dt and AMPA:NMDA ratio, comparable to that of local stereotactic *Scn2a*-rAAV-CRISPRa treatment directly into the mPFC (**Fig. 3C-D**). Together, these data suggest the feasibility of systemic administration of rAAV-CRISPRa to rescue *Scn2a* haploinsufficiency-associated electrophysiological defects.

**Figure 3.**
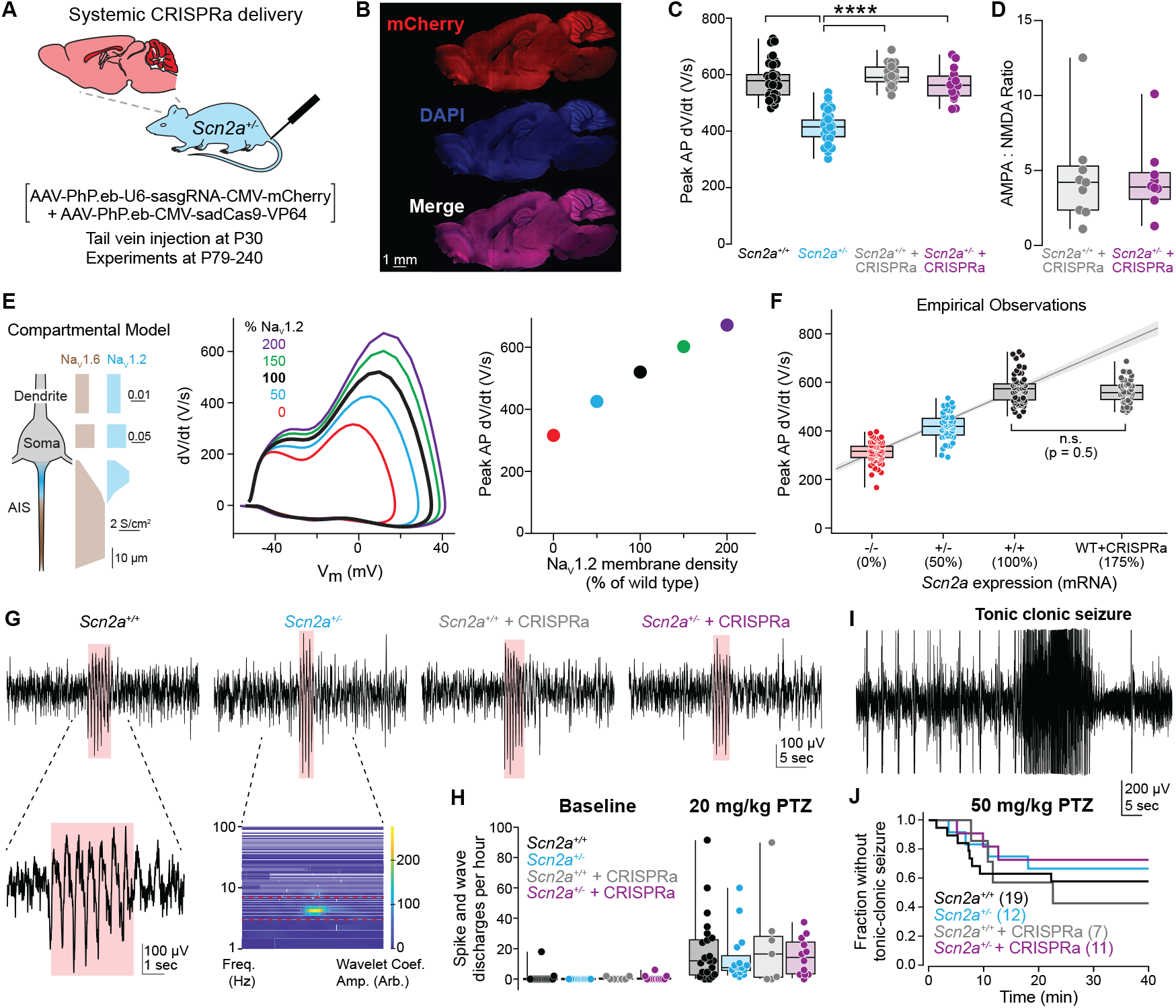
Systemic delivery of CRISPRa rescues electrophysiological deficits without increasing seizure burden. **A:** Schematic of intravenous tail vein delivery of *Scn2a*-rAAV-CRISPRa-PhP.eb and ubiquitous neuronal mCherry expression. Viruses injected described in brackets. **B:** Sagittal brain section from P80 *Scn2a*^+/-^ + CRISPRa mouse following tail vein injection of *Scn2a*-rAAV-CRISPRa-PhP.eb at P30-40 immunostained with antibodies against mCherry and DAPI. **C:** Quantification of peak AP dV/dt from P82-241 *Scn2a*^+/+^ (black), *Scn2a*^+/-^ (cyan), and tail vein *Scn2a*-rAAV-CRISPRa-PhP.eb injected *Scn2a*^+/+^ + CRISPRa (gray) and *Scn2a*^+/-^ + CRISPRa (purple) neurons. Circles represent individual neurons. *Scn2a*^+/+^: 578.2 ± 9.3, n = 44 cells; *Scn2a*^+/-^: 413.8 ± 8.8, n = 37 cells; *Scn2a*^+/+^ + CRISPRa: 590.0 ± 8.4, n = 23 cells; *Scn2a*^+/-^ + CRISPRa: 561.2 ± 11.5 cells, n = 20 cells. *Scn2a*^+/+^ vs. *Scn2a*^+/-^: ****p < 0.0001, *Scn2a*^+/-^ vs. *Scn2a*^+/+^ + CRISPRa: ****p < 0.0001, *Scn2a*^+/-^ vs. *Scn2a*^+/-^ + CRISPRa: ****p < 0.0001. Holm-Šídák multiple comparisons test. **D:** AMPA:NMDA ratio from P82-241 *Scn2a*-rAAV-CRISPRa-PhP.eb treated *Scn2a*^+/+^ and *Scn2a*^+/-^ mice. *Scn2a*^+/+^ + CRISPRa: 4.3 ± 0.9, n = 11 cells; *Scn2a*^+/-^ + CRISPRa: 4.6 ± 3.1, n = 13 cells. p = 0.7. Mann-Whitney test. **E:** Left: Schematic of predicted distribution of NaV1.2 and NaV1.6 channels across the axon initial segment (AIS), soma, and proximal apical dendrite. Middle: Phase-plane plots from computational model of peak dV/dt across varying amounts of NaV1.2 channel densities in plasma membrane. Right: Predicted peak AP dV/dt as a function of NaV1.2 expression levels predicted by computational modeling. **F:** Empirical measurements of peak AP dV/dt in *Scn2a*^+/+^, *Scn2a*^+/-^, and *Scn2a*^-/-^ neurons compared to *Scn2a*^+/+^ + CRISPRa. *Scn2a*^+/+^, *Scn2a*^+/-^, and *Scn2a*^-/-^ values are from Spratt et al. 2021. *Scn2a*^+/+^ + CRISPRa are merged local and tail vein peak dV/dt values. Data placed at 175% of WT based on PCR upregulation (Fig. 2C). *Scn2a*^+/+^ vs. *Scn2a*^+/+^ + CRISPRa: no significance (p = 0.5). Mann-Whitney test. **G:** Recordings of (SWDs) following 20 mg/kg PTZ administration in P79-148 *Scn2a*^+/+^, *Scn2a*^+/-^, and *Scn2a*-rAAV-CRISPRa-PhP.eb treated *Scn2a*^+/+^ and *Scn2a*^+/-^ mice. Insets highlight SWD in *Scn2a*^*+/+*^ mouse and spectrogram with high wavelet coefficient amplitude in the 3-5 Hz band range (red dashed region) in *Scn2a*^*+/-*^ example. **H:** Left: Quantification of SWDs detected at baseline. *Scn2a*^+/+^ (black): 0.8 ± 0.8, n = 24 mice; *Scn2a*^+/-^ (cyan): 0 0, n = 16 mice; *Scn2a*^+/+^ + CRISPRa (gray): 0.6 ± 0.4, n = 7 mice; *Scn2a*^+/-^ + CRISPRa (purple): 0.8 ± 0.5 mice, n = 12. No significant differences. Holm-Šídák multiple comparisons test. Right: SWDs following low dose treatment (20 mg/kg) of pentylenetetrazole (PTZ). *Scn2a*^+/+^: 18.2 ± 5.5, n = 24 mice; *Scn2a*^+/-^: 14.1 ± 4.1, n = 16 mice; *Scn2a*^+/+^ + CRISPRa: 23.6 ± 12.1, n = 7 mice; *Scn2a*^+/-^ + CRISPRa: 14.9 ± 3.6, n = 12 mice. No significant differences. Holm-Šídák multiple comparisons test. **I:** Example of a tonic clonic seizure induced with a high dose (50 mg/kg) administration of PTZ in P100-167 mice. **J:** Survival curves following high dose (50 mg/kg) treatment of PTZ over 40 minutes of EEG recordings. P = 0.69, Mantel log-rank test across comparisons.

### *Scn2a* upregulation in WT mice does not induce phenotypic abnormalities

Gain-of-function missense variants in *SCN2A* are associated with multiple forms of epilepsy (Reynolds et al., 2020). Thus, a major concern with our CRISPRa approach to rescue *SCN2A* haploinsufficiency is overexpression beyond normal physiological levels, as this may result in hyperexcitability, similar to that observed with gain-of-function missense variants. To assess this, we administered intracranially CRISPRa in WT mice and assessed their cellular excitability. Intracranial *Scn2a*-rAAV-CRISPRa injection into mPFC upregulated *Scn2a* mRNA ∼1.75-fold in WT mice (**Fig. 2C**).

Compartmental models predicted that such increases in mRNA, if translated into Na_V_1.2 channels that incorporate into neuronal membranes, should increase peak dV/dt of APs (**Fig. 3E**). Indeed, AP velocity in such models was proportional to Na_V_1.2 membrane density from 0 to 200% of normal levels. In empirical studies, we observed a similar linear relationship between AP peak dV/dt and *Scn2a* copy number between 0 and 100% of normal levels; however, *Scn2a*-rAAV-CRISPRa-treated *Scn2a*^*+/+*^ cells had peak dV/dt values that were no different than untreated controls (**Fig. 3F**). This suggests that maximal Na_V_1.2 membrane expression is regulated, perhaps by auxiliary subunits or scaffolding partners, and cannot exceed WT levels (Hull and Isom, 2018).

### *Scn2a* upregulation does not induce seizures

While single-cell electrophysiology suggests that overexpression of *Scn2a* beyond WT levels does not result in intrinsic hyperexcitability in neocortical neurons, upregulation of this gene in other cell types could result in network-level epileptogenic effects. Therefore, we assessed overall brain excitability with electroencephalography (EEG) in awake behaving mice, both heterozygous and WT, that received systemic PhP. eb-based CRT. EEG was assessed in naïve mice and following two different doses of the convulsant pentylenetetrazole (PTZ, 20 mg/kg or 50 mg/kg). Epilepsy-associated spike and wave discharges (SWD), which are present at low frequency in EEG recordings of naïve C57BL/6 mice and increase in frequency in response to PTZ (Purtell et al., 2018), were no different in frequency across cohorts of CRT-treated and untreated *Scn2a*^*+/+*^ and *Scn2a*^*+/-*^ mice in baseline conditions or following 20 mg/kg PTZ administration (**Fig. 3G-H**). At the higher PTZ dose (50 mg/kg), some animals experienced behavioral arrest followed by seizures of increasing severity, including tonic-clonic seizures. The prevalence of these events was not different across cohorts (**Fig. 3I-J**). These data suggest that *Scn2a*-rAAV-CRISPRa is unlikely to increase seizure burden, regardless of overall *Scn2a* expression levels.

### Rescue of electrophysiological deficits in human *SCN2A+/-* excitatory neurons

To further investigate the translational potential of this approach, we tested the ability of *SCN2A*-rAAV-CRISPRa to rescue the electrophysiological phenotypes of *SCN2A*^+/-^ neurons differentiated from human embryonic stem cells (hESCs). sgRNA constructs that target the human *SCN2A* promoter were designed in a similar strategy to those for mouse and screened for upregulation of *SCN2A* in a human neuroblastoma cell line (SH-SY5Y). One sgRNA, out of the ten tested, increased *SCN2A* mRNA by 1.5- and 1.3-fold following transient co-transfection with dCas9-VP64 or transduced via a rAAV-DJ serotype, respectively (**Fig. S6A-B**). *SCN2A*^*+/+*^ and *SCN2A*^*+/-*^ hESCs (Lu et al., 2019) were differentiated into excitatory neurons using the SMADi STEMDiff forebrain differentiation and maturation protocol (**Fig. 4A**) (Ruden et al., 2021). Immunostaining of cells at DIV 65 with MAP2 and GFAP suggested efficient neuronal differentiation alongside supporting glial cells (**Fig. 4B**).

**Figure 4.**
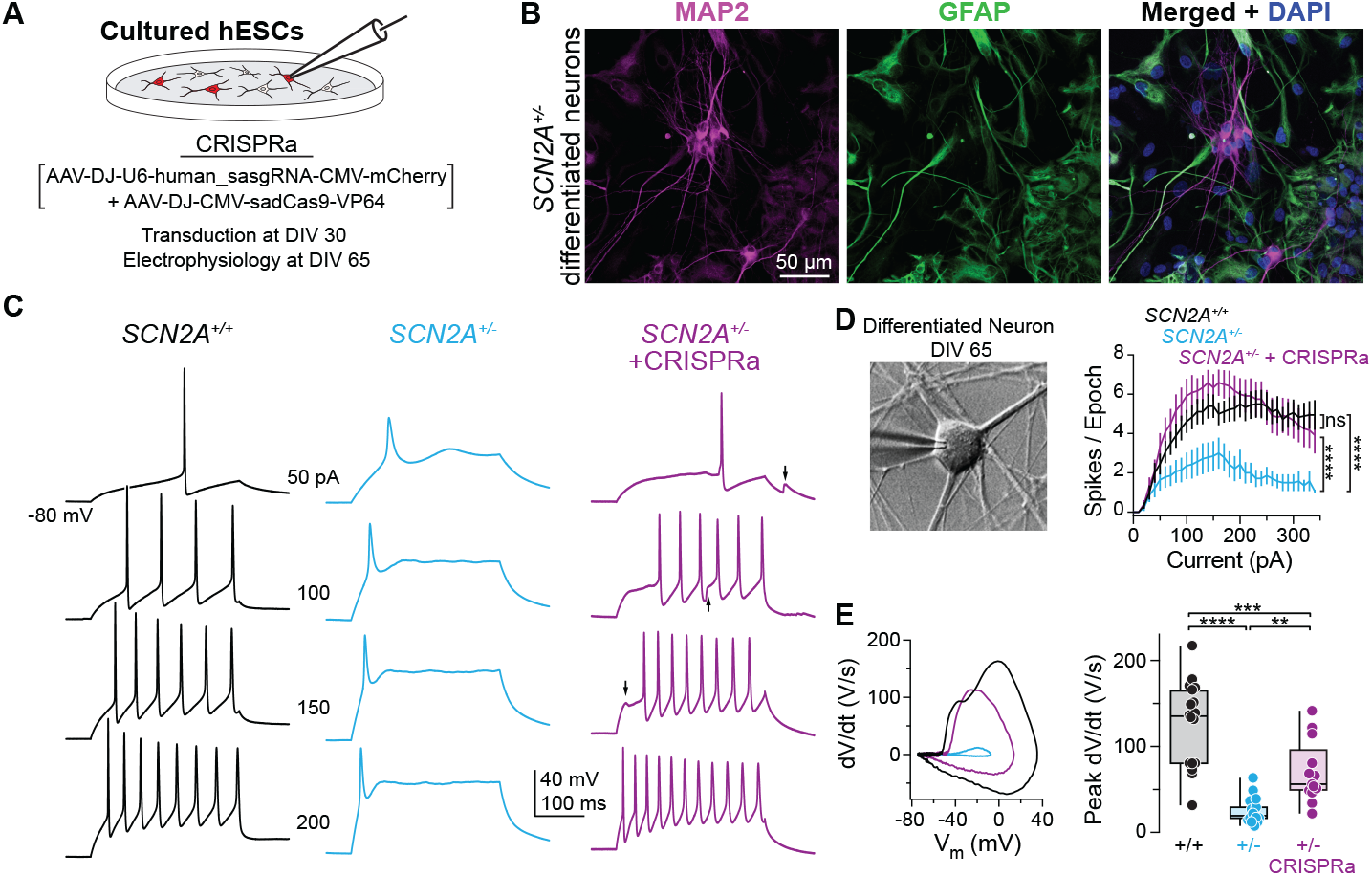
*SCN2A-*rAAV-CRISPRa rescues spiking properties of *SCN2A*^+/-^ human embryonic stem cell-derived neurons. **A:** Schematic of electrophysiological recording of CRISPRa-treated differentiated hESC-derived neurons. Cells were transduced at DIV 30, and experiments were performed at DIV 65-66. **B:** Differentiated *SCN2A*^+/-^ hESC-derived neurons at DIV 65 immunostained with antibodies against the neuronal marker MAP2, the glial marker GFAP, and DAPI. **C:** APs generated from a range of current amplitudes (0-340 pA, 10 pA intervals, 300 ms) in DIV 65-66 *SCN2A*^+/+^ (black), *SCN2A*^+/-^ (cyan), and *SCN2A*-rAAV-CRISPRa treated *SCN2A*^+/-^ (purple) hESC-derived neurons. Arrows highlight spontaneous excitatory postsynaptic potentials (EPSPs) that occurred within recording epoch. **D:** Left: DIC image of *SCN2A*^+/-^ hESC-derived neuron and recording electrode. Right: AP (spikes) per 300 ms stimulation epoch versus current injection amplitude. *SCN2A*^+/+^ vs. *SCN2A*^+/-^: ****p < 0.0001. *SCN2A*^+/-^ vs. *SCN2A*^+/-^ + CRISPRa ****p < 0.0001. Kruskal-Wallis test. **E:** Representative phase-plane plots and quantification of peak dV/dt in *SCN2A*^+/+^, *SCN2A*^*+/-*^, and *SCN2A*-rAAV CRISPRa treated *SCN2A*^+/-^ hESC-derived neurons. *SCN2A*^+/+^ 123.8 ± 11.7, n = 18 cells; *SCN2A*^+/-^ 24.7 ± 3.4, n = 18 cells; *SCN2A*^+/-^ + CRISPRa 69.7 ± 9.9, n = 13 cells. *SCN2A*^+/+^ vs. *SCN2A*^+/-^: ****p < 0.0001. *SCN2A*^+/+^ vs. *SCN2A*^+/-^ + CRISPRa: ***p = 0.0004. *SCN2A*^+/-^ vs. *SCN2A*^+/-^ + CRISPRa: ** p = 0.0014. Holm-Šídák multiple comparisons test.

Early in life, both human and rodent excitatory pyramidal cells rely on Na_V_1.2 expression in the AIS to support AP electrogenesis (Gazina et al., 2015). During this period, *Scn2a* haploinsufficiency suppresses AP output (Spratt et al., 2019). This developmental period occurs over one year in life in humans (Workman et al., 2013). As such, CRT can be evaluated in hESCs by examining both AP waveform and AP output. At DIV 65-66, *SCN2A*^*+/+*^ neurons were mature enough to support repetitive AP activity (**Fig. 4C**). By contrast, AP electrogenesis was markedly blunted in *SCN2A*^*+/-*^ neurons, with few evoked APs that had very low peak dV/dt values compared to interleaved controls (**Fig. 4D-E**). Furthermore, the axon initial segments of these *SCN2A*^*+/-*^ neurons were ∼19% longer than those from *SCN2A*^*+/+*^ counterparts (**Fig. S7A-B**). This is consistent with structural plasticity of this compartment associated with reduced neuronal excitability that can be observed following changes in activity and in response to disease-associated alterations in Na_V_ function (Leterrier, 2018; Tidball et al., 2020). *SCN2A*^*+/-*^ neurons treated with *SCN2A*-rAAV-CRISPRa at DIV 30 had mRNA levels that were comparable to WT levels (**Fig. S5**) and at DIV 65-66 exhibited a full recovery in AP output with a concomitant increase in peak dV/dt (**Fig. 4C-E**). Moreover, CRISPRa-infected *SCN2A*^*+/-*^ initial segments were 19% shorter than those of uninfected *SCN2A*^*+/-*^ neurons visualized in the same preparation (**Fig. S7C-D**). In conclusion, these data demonstrate that CRISPRa can provide a potential therapeutic benefit for *SCN2A* haploinsufficiency in human neuronal cell types, rescuing both electrical and structural aspects of neuronal excitability.

## DISCUSSION

One hundred and sixty-one high-confidence ASD risk genes are thought to cause ASD due to haploinsufficiency, with *SCN2A* haploinsufficiency being one of the most significant genetically defined subsets of ASD (Karczewski et al., 2020). Here, we showed how a CRISPRa-based CRT approach could be used to rescue the electrophysiological deficits associated with *SCN2A* haploinsufficiency. First, using a *Cre*-induced rescue of *Scn2a* haploinsufficiency in adolescent mice, we showed the feasibility of this approach to provide a potential therapeutic during adolescence. Next, by direct stereotaxic mPFC and intravenous tail vein injections of *Scn2a-* rAAV-CRISPRa, we upregulated *Scn2a* in adolescent *Scn2a*^+/-^ mice and demonstrated rescue of multiple features of intrinsic excitability, including AP electrogenesis and associated activation of dendritic calcium channels. These intrinsic changes were associated with rescue of synaptic properties that are typical of immature neurons, including low AMPA:NMDA ratio and spines with a more filipodial morphology. Of note, these synaptic features were rescued with CRISPRa administration in early adolescence, suggesting that both *Scn2a* haploinsufficiency interrupts normal circuit refinement and that these processes can be reinitiated if normal levels of Na_V_1.2 are restored. Together, these results suggest that CRT is a viable therapeutic approach for *SCN2A* haploinsufficiency.

In the case of sodium channel haploinsufficiency, CRT also overcomes two of the main limitations of traditional gene replacement therapy, rAAV packaging capacity and ectopic expression. Sodium channels are long genes whose cDNA exceed the rAAV payload capacity. CRT overcomes this by packaging CRISPRa components that target the gene’s endogenous regulatory elements to upregulate its expression. In addition, results from a transgenic-based CRISPRa suggest that upregulation occurs only in the tissue/cell type where the targeted regulatory element is active (Matharu et al., 2019), providing additional specificity to the rAAV serotype or promoter used to drive expression. In particular, this could be extremely beneficial for sodium channels that have similar protein structures yet have distinct functions in skeletal, cardiac, central, and peripheral nervous systems, which makes it difficult to target specific proteoforms pharmacologically without off-target effects (Johnson et al., 2022; Waszkielewicz et al., 2013). An additional advantage is that CRT uses a nuclease-deficient DNA targeting molecule that does not edit the DNA, and thus off-target effects will not lead to ‘DNA scars’. The use of a CRISPR-based CRT approach could lead to immunogenicity due to the dCas9. However, there are many efforts to engineer Cas proteins with reduced immunogenicity by epitope masking (Ferdosi et al., 2019; Mehta and Merkel, 2020), or use of alternate Cas proteins, such as those from non-pathogenic bacteria that have not been exposed to humans (Matharu and Ahituv, 2020). Alternatively, other DNA targeting molecules like zinc fingers or Transcription Activator-Like Effector Nucleases (TALENS), that should be less immunogenic and smaller in size (Gaj et al., 2016), could be used instead.

A major goal of CRT is to achieve physiological levels of the modulated gene that provide therapeutic benefit. Obtaining levels that are too high could also have deleterious physiological impacts whereas levels too low may not be within the therapeutic window of efficacy. For *SCN2A*, this is of particular importance as gain-of-function variants are associated with epileptic encephalopathies (Spratt et al., 2021; Wolff et al., 2017). Here, we intentionally used VP64 as the transcriptional activator to avoid excess Na_V_1.2 production, as it has modest upregulating potential (Chavez et al., 2016, p. 201). Furthermore, we also assessed whether *Scn2a* over-expression via CRISPRa above normal physiological conditions affected brain function by upregulating *Scn2a* mRNA in WT mice. Heightened levels of *Scn2a* mRNA did not correspond to elevated intrinsic or network neuronal hyperexcitability, nor did they lead to any change in seizure burden in naïve or convulsant-treated mice. We speculate that this could be due to a potential ceiling on total Na_V_1.2 membrane density, perhaps imposed by ancillary subunits or scaffolding partners whose expression remains unchanged (Colasante et al., 2020; Hull and Isom, 2018; Isom and Knupp, 2021). Furthermore, we observed no expression or electrophysiological effects that would indicate that CRT altered the expression of other CNS sodium channel genes within the *Scn2a* TAD region (*Scn1a, Scn3a*) or in other sodium channels expressed in pyramidal cells (*Scn8a*, which is critical for AP threshold).

Treatments for NDDs would likely be most beneficial when administered early, ideally before symptom onset (Marín, 2016). Nevertheless, brain development is a dynamic process that spans multiple decades of life, and interventions administered even later in life could have some therapeutic benefit (Levy and Barak, 2021). A recent example is an antisense oligonucleotide therapy for Angelman Syndrome that improved communication and motor skills and reduced epileptiform discharges when administered to 4–17-year-old children (Markati et al., 2021), despite work in rodent models suggesting that therapies should ideally be administered prenatally (Silva-Santos et al., 2015; Wolter et al., 2020). *Scn2a* haploinsufficiency in mice results in life-long, cell-autonomous impairments in neocortical pyramidal cell dendritic excitability and synapse function (Spratt et al., 2019). Here, we find that these impairments can be rescued with adolescent reactivation of a loss-of-function allele (**Fig. 1**) or by CRT-based over-expression via the residual, functional allele (**Fig. 2-3**), suggesting that rescue of normal dendritic excitability can restore many aspects of neuronal function. Leveraging other model systems, including non-human primates, will likely be critical for assessing proper therapeutic developmental windows and safety profiles. This will be especially important since behavioral abnormalities in *Scn2a* haploinsufficient mice are difficult to reliably observe across studies, which is consistent with many other heterozygote models of ASD-related genes (Jiang and Ehlers, 2013; Shin et al., 2019; Spratt et al., 2019; Tatsukawa et al., 2019; Zhou et al., 2019). Furthermore, several aspects of this approach will need to be optimized, including safety assessments of rAAV components, off-target effects, the use of non-human primates to assess efficient delivery, upregulation levels, cytotoxicity, immunogenicity and other effects (Goertsen et al., 2022). In addition, it will be vital to confirm that the *SCN2A* variant to be treated causes a non-functional transcript, as this current approach upregulates both alleles.

In summary, this study shows how CRT using CRISPRa can rescue a major class of mutations in a significant ASD risk gene. CRT is a customizable platform technology that can modify gene expression without directly editing the genome and can be tailored to rescue other disorders of haploinsufficiency, including other NDDs. The application of CRT to treat *SCN2A* haploinsufficiency leverages a growing understanding of the genetic etiology of ASD and demonstrates a potential path forward for treating complex behaviorally defined conditions like ASD through modern gene therapy approaches.

## MATERIALS AND METHODS

### CRISPRa *In Vitro* Optimization

Ten sgRNAs targeting the mouse *Scn2a* or human *SCN2A* promoters were designed using the Broad Institute’s GPP sgRNA Design Tool (Genetic Perturbation Platform, Broad Institute). These guides were individually cloned into pAAV-U6-sasgRNA-CMV-mCherry-WPREpA at the *BstX*I and *Xho*I restriction enzyme sites using the In-Fusion HD cloning kit (Clontech). rAAV vectors were generated using similar plasmids and cloning methods as was referenced in (Matharu et al., 2019). Mouse sgRNAs were tested in the neuroblastoma cell line Neuro-2a (ATCC CCL-131). Cells were grown in Dulbecco’s Modified Eagle’s Medium (DMEM) with 10% FBS and 1% Penicillin-Streptomycin, and SH-SY5Y cells were grown in Eagle’s Minimum Essential Medium (EMEM) with 10% FBS and 1% Penicillin-Streptomycin following ATCC guidelines. Cells were transiently co-transfected with individual sgRNA cloned into pAAV-U6-sasgRNA-CMV-mCherry-WPREpA along with pCMV-sadCas9-VP64 for 48 hours using Opti-MEM Reduced Serum Medium (Thermo Fisher) and X-tremeGENE HP (Sigma-Aldrich). RNA was isolated using the RNeasy Mini Kit (Qiagen) following the manufacturer’s protocol. cDNA was synthesized using SuperScript III First-Strand Synthesis System (Invitrogen) and qPCR was conducted using SsoFast EvaGreen Supermix (Bio-Rad) and analyzed using the ΔΔCT methods comparing to a no-sgRNA transfection and normalized to Actb as a housekeeping gene. rAAVs were produced at the Stanford Gene Vector and Viral Core (see **Table S3** for genomic titers). RNA-seq, using four biological replicates per each condition, was carried out on a NovaSeq 6000 generating 150 base pair end reads. Sequences were mapped to mm10 with STAR 2.5.1b (Dobin et al., 2013) and counts were computed using featureCounts with gtf provided from genecode (gencode.vM25.primary_assembly.annotation.gtf). Differential expression was characterized using DESeq2 R version 4.1.2 (2021-11-01) (Love et al., 2014). All RNA-seq data is available on the NCBI Gene Expression Omnibus as Bioproject GSE193605.

### Mouse Generation, Husbandry, and Genotyping

All mouse work was approved by the UCSF Institutional Animal Care and Use Committee. All mice were maintained on a C57BL/6J background on a 12:12 light-dark cycle (light on at 6 AM) with ad libitum food and water. Genotyping was performed via tail clips using the KAPA Mouse Genotyping Kit (Roche #07961766001). See **Table S2** for primer sequences. *Scn2a*^*+/-*^ were as described in (Planells-Cases et al., 2000), and conditional knockout (Scn2a^+/fl^) were as described in (Spratt et al., 2019). *Scn2a*^*+/KI*^ mice were created at Cyagen Biosciences by inverting exons 3-5 of the *Scn2a* gene and inserting a SA-2A-eGFP-polyA cassette upstream of the inverted exons. The inverted exons and the SA-2A-eGFP-polyA cassette were flanked with loxP and Lox2272 sites to ensure the excision of the SA-2A-eGFP-polyA cassette and the reversion of exons 3-5 to the correct orientation following Cre recombinase induced recombination. A targeting vector containing the inverted exons, SA-2A-eGFP-polyA cassette, recombination sites, selection markers, and homology arms was assembled from mouse genomic fragments amplified from BAC clone RP23-332C13 and RP23-55C23. The targeting vector was linearized by restriction enzyme digestion and transfected into C57BL/6 embryonic stem cells via electroporation, and successfully transfected cells were identified by drug selection, PCR verification, and Southern blot confirmation. Confirmed clones were introduced into host embryos and transferred to surrogate mothers. Chimerism in the resulting pups was identified via coat color. F0 male chimeras were bred with C57BL/6J females to generate F1 heterozygous mutants that were identified by PCR.

### *In Vivo* AAV Administration

For local AAV administration, mice P30-P40 of age were kept under live anesthetic isoflurane at 0.5-2.0% and mounted onto the stereotaxic machine (Kopf 1900). 500 nL of rAAV at a 1:1 ratio of sgRNA and sadCas9-VP64 was injected into the mPFC at stereotaxic coordinates [mm]: anterior-posterior [AP], +1.7, mediolateral [ML] −0.35; dorsoventral [DV]: −2.6 at a viral infusion rate of 0.1 μl min−1. For systemic AAV administration, mice age P30-P40 were kept on a 37° C warm pad and harnessed using a brass mouse restrainer (SAI Infusion Tech). Lateral tail vein injections were carried out with rAAV at a 1:1 ratio of sgRNA and sadCas9-VP64 (1×10^11 vg/mouse) suspended in 200 uL saline using a 30G needle. For either injection, mice were used for subsequent experiments four weeks post-injection. Tail vein injected animals used for EEG recording experiments, were subsequently divided across experiments for electrophysiological recordings, qPCR, and immunofluorescence.

### *In Vivo* Dissections

Four weeks post-injection, mouse brains were removed and 250 μm thick coronal slices containing the mPFC were dissected in artificial cerebrospinal solution containing (in mM): 87 NaCl, 25 NaHCO_3_, 25 glucose, 75 sucrose, 2.5 KCl, 1.25 NaH_2_PO_4_, 0.5 CaCl_2_ and 7 MgCl_2_; bubbled with 5%CO_2_/95%O_2_; 4°C. Fluorescence mCherry expression in the mPFC of the injected hemisphere was validated using a fluorescent stereomicroscope (Nikon SMZ1500). For mRNA expression analyses, both the injected (mCherry-positive) and uninjected hemisphere (mCherry-negative) were dissected using a sterile miltex disposable punch biopsy (Medline MIL3332P25) and flash frozen in RLT lysis buffer (Qiagen). RNA was extracted using the RNAEasy Micro Kit (Qiagen) following the manufacturer’s protocol. cDNA and downstream qPCR were conducted using SuperScript III (Invitrogen) and SsoFast EvaGreen Supermix (Bio-Rad) on a QuantStudio 6 Flex Real Time PCR system (Applied Biosystems). qPCR results comparing the injected and uninjected hemispheres were analyzed using the ΔΔCT methods and normalized to *Actb*, a housekeeping gene.

### *Ex Vivo* Electrophysiology and Two-photon Imaging

All *ex vivo* electrophysiology and two-photon imaging were acquired and performed using the same methods as previously described (Spratt et al., 2021, 2019). Mice were anesthetized using isoflurane and 250 μm coronal slices were prepared. Cutting solution contained (in mM): 87 NaCl, 25 NaHCO_3_, 25 glucose, 75 sucrose, 2.5 KCl, 1.25 NaH_2_PO_4_, 0.5 CaCl_2_ and 7 MgCl_2_; bubbled with 5%CO_2_/95%O_2_; 4°C. Following cutting, slices were either incubated in the same solution or in the recording solution for 30 min at 33 °C, then at room temperature until recording. Recording solution contained (in mM): 125 NaCl, 2.5 KCl, 2 CaCl_2_, 1 MgCl_2_, 25 NaHCO_3_, 1.25 NaH_2_PO_4_, 25 glucose; bubbled with 5%CO_2_/95%O_2_; 32–34°C, ∼310 mOsm. Neurons were visualized with differential interference contrast (DIC) optics for conventional visually guided whole-cell recording, Dodt contrast imaging, or with 2-photon-guided imaging of reporter-driven mCherry fluorescence overlaid on an image of the slice (scanning DIC). For current-clamp recordings, patch electrodes (Schott 8250 glass, 3–4 MΩ tip resistance) were filled with a solution containing (in mM): 113 K-Gluconate, 9 HEPES, 4.5 MgCl_2_, 0.1 EGTA, 14 Tris_2_-phosphocreatine, 4 Na_2_-ATP, 0.3 tris-GTP; ∼290 mOsm, pH: 7.2–7.25. For Ca^2+^ imaging, EGTA was replaced with 250 μM Fluo-5F and 20 μM Alexa 594. For voltage-clamp recordings of and synaptic activity, internal solution contained (in mM): 110 CsMeSO_3_, 40 HEPES, 1 KCl, 4 NaCl, 4 Mg-ATP, 10 Na-phosphocreatine, 0.4 Na_2_-GTP, 0.1 EGTA; ∼290 mOsm, pH: 7.22. All data were corrected for measured junction potentials of 12 and 11 mV in K- and Cs-based internals, respectively.

Electrophysiological data were acquired using Multiclamp 700A or 700B amplifiers (Molecular Devices) via custom routines in IgorPro (Wavemetrics). For measurements of action potential waveform, data were acquired at 50 kHz and filtered at 20 kHz. For all other measurements, data were acquired at 10–20 kHz and filtered at 3–10 kHz. For current-clamp recordings, pipette capacitance was compensated by 50% of the fast capacitance measured under gigaohm seal conditions in voltage-clamp prior to establishing a whole-cell configuration, and the bridge was balanced. For voltage-clamp recordings, pipette capacitance was compensated completely, and series resistance was compensated 50%. Series resistance was <15 MΩ in all recordings. Experiments were omitted if input resistance changed by > ±15%.

Two-photon laser scanning microscopy (2PLSM) was performed as previously described (Spratt et al., 2019). A two-photon source (Coherent Ultra II) was tuned to 810 nm for morphology and calcium imaging. Epi- and transfluorescence signals were captured either through a 40×, 0.8 NA objective for calcium imaging or a 60×, 1.0 NA objective for spine morphology imaging, paired with a 1.4 NA oil immersion condenser (Olympus). Fluorescence was split into red and green channels using dichroic mirrors and band-pass filters (575 DCXR, ET525/70m-2p, ET620/60m-2p, Chroma). Green fluorescence (Fluo-5F) was captured with 10770–40 photomultiplier tubes selected for high quantum efficiency and low dark counts (PMTs, Hamamatsu). Red fluorescence (Alexa 594) was captured with R9110 PMTs. Data were collected in linescan mode (2–2.4 ms/line, including mirror flyback). For calcium imaging, data were presented as averages of 10–20 events per site and expressed as Δ(G/R)/(G/R) max*100, where (G/R)max was the maximal fluorescence in saturating Ca2+ (2 mM) (Yasuda et al., 2004). Action potential backpropagation experiments were performed in 25 μM picrotoxin, 10 μM NBQX and 10 μM R-CPP.

### EEG Implant Surgeries

Mice were anesthetized with isoflurane and placed on a stereotaxic apparatus. Screws with wire leads were implanted in 2-3 turns into five burr holes at (stereotaxic coordinates relative to bregma [mm]) PFC: 1.7 anterior-posterior (AP), −0.3 mediolateral (ML); S1: −1.8 AP, 2.5 ML; reference and ground: −5 AP, 0.9 ML (Pinnacle Technology). A head mount was then attached to the wire leads and secured in adhesive dental cement. Mice were given at least one week for post-surgery recovery prior to EEG recordings.

### EEG Recordings

EEG recordings were performed using the Sirenia Acquisition Software (Pinnacle Technology) at 2 kHz sampling frequency with simultaneous video capture. Pentylenetetrazol (PTZ) administration of low (20 mg/kg) and high (50 mg/kg) doses were performed in two separate sessions at least two weeks apart to minimize the kindling effect. PTZ solutions were prepared fresh at the start of every recording day. For each session, baseline activity was recorded for 30 minutes after which the mice were injected with PTZ via intraperitoneal injection and continued recording for 40 minutes. Recordings were performed between 10 AM and 6 PM to control for the circadian light-dark cycle.

### EEG Analysis

All analyses were performed offline using custom software written in Python. Preprocessing was first performed on baseline and low dose PTZ recordings including a bandpass filter between 1-100 Hz before downsampling to 200 Hz. Spectral power was computed using the continuous wavelet transform (CWT) with a complex Morlet wavelet. The recording was segmented into 125 ms windows and classified as part of an SWD based on the amplitude of spikes and the spectral power. The band power of interest was calculated as the 3-7 Hz band subtracted from the 0-1 Hz band to eliminate non-SWD frequencies. Spikes in the EEG signal were detected based on an amplitude threshold of 3x the root mean square of baseline and a power threshold of 3x the band power in the first minute of baseline. SWDs were considered events with durations lasting at least 1 sec and verified by manual inspection. Power spectral density was calculated using the Welch’s method on z-score normalized EEG signal over the entire baseline or high dose PTZ session.

Seizure scoring for high dose PTZ recordings were measured from simultaneous video and EEG monitoring. Behavioral classification of increasing seizure severity was based on the Racine scale and a previous report of PTZ-induced seizures (Van Erum et al., 2019 & Miyamoto et al., 2019). Latency to seizure activity was considered the time from PTZ administration to the onset of the first seizure. Behavioral arrest was defined as a sudden immobilization accompanied by SWDs. A myoclonic jerk was defined as a neck or body twitch accompanied by a single sharp spike in the EEG signal. A tonic-clonic seizure was defined as clonic convulsions leading to wild jumping also with high amplitude spiking in the EEG signal. The animal was considered deceased when the tonic-clonic seizure led to tonic extension of hind limbs followed by loss of body tone and close to flat EEG signal.

### Computational Compartmental Modeling

A pyramidal cell compartmental model was implemented in the NEURON environment (v7.7) based on the Blue Brain Project thick-tufted layer 5b pyramidal cell (TTPC1) model used in our previous study (Ben-Shalom et al., 2017; Markram et al., 2015; Spratt et al., 2021). The TTPC1 model was adjusted to include an AIS, and the original Na channels in the TTPC1 model were replaced with Na_V_1.2 and Na_V_1.6 channels in compartments with densities as described previously (Spratt et al., 2021).

### hESC Differentiation, Maturation, and Electrophysiology

Wildtype and *SCN2A*^+/-^ HUES66 (NIH Registration #0057) hESC cell lines were obtained from the Harvard Stem Cell Institute and plated on Matrigel (Corning) coated standard tissue culture plates maintained in mTESR (STEMcell technologies). NPCs hESCs were differentiated following the manufacturer’s instructions using the STEMDiff SMADi Embryoid Body Neural Induction protocol (STEMcell Tech) following a similar strategy documented in (Ruden et al., 2021). Neural progenitor cellsNPCs were further differentiated into neuronal forebrain precursors using the STEMdiff ForeBrain Neuron Differentiation protocol (Document #10000005464, Stemcell Technologies, Vancouver, CanadaSTEMcell Tech), and neuronal precursors were matured into forebrain neurons using the STEMdiff Forebrain Maturation Kit with BrainPhys (STEMCell Tech). Neuronal precursors were maturated on Poly-L-Ornithine Laminin coated German glass coverslips (Neuvitro GG-25-1.5-Laminin) for 65 days. Whole-cell current-clamp recordings were done as in mouse acute slices using identical solutions.

### Immunofluorescence of Fixed Cells or Tissue

For hESCs, differentiated neurons were fixed at DIV 65 with 4% (paraformaldehyde) PFA and 4% sucrose and blocked with 10% Normal Goat Serum in phosphate buffered saline (PBS) in 0.2% Triton-X (PBST). All coverslips within a set of experiments, including those used previously that day for electrophysiology, were fixed within 1 hr of each other to minimize developmental differences across coverslips. Following fixation, cells were incubated with primary antibodies against MAP2 (Invitrogen PA1-10005) at 1:5000 and Ankyrin-G (Neuromab 75-146) at 1:1000 overnight at 4 degrees. Cells were then incubated with the secondary antibodies Goat anti-Chicken IgY (H+L) Cross-Adsorbed Secondary Antibody, Alexa Fluor Plus 647 (Invitrogen A32933) at 1:500 and Goat anti-Mouse IgG (H+L) Highly Cross Adsorbed Secondary Antibody, Alexa Fluor 488 (Thermo Fisher A-11029) at 1:500. They were then mounted on coverslips with ProLong(tm) Diamond Antifade Mountant with DAPI (Thermo Fisher). Images were captured with a 40x 1.4NA objective using an Olympus Fluoview FV3000 confocal microscope. AIS length was measured from maximum intensity projections generated from z-stacks that contained the entire bounds of the neuron based on ankyrin-G and MAP2 immunofluorescence. Only initial segments with clearly detectable start and end points were quantified using the segmented line tool in FIJI (ImageJ). Start and end points were determined with a threshold of 50% peak ankyrin-G fluorescence intensity using plot profiles (pixel intensity/distance) in FIJI.

For whole-brain mouse immunohistochemistry, mice were perfused with 4% PFA in PBS and brains were removed and sectioned coronally or sagittally on vibratome (Leica VT1000) at 50-75 μm and mounted on slides with ProLong(tm) Gold Antifade Mountant with DAPI. For imaging of *Scn2a*^*+/KI*^ sections, tissue was permeabilized, blocked, and immunostained for GFP (anti-GFP AlexaFluor 488, 1:500, ThermoFisher A-21311) and parvalbumin (PV27, 1:500, Swant; AlexaFluor 564 secondary, 1:500, ThermoFisher). Sections were imaged either with an Olympus Fluoview FV3000 (coronal sections) or Keyence BZ-X widefield microscope (sagittal sections).

### Quantification and Statistical Analysis

Unless otherwise noted, all data are shown in figures as box plots with median and quartiles and min. and max. tails with individual data points overlaid. Data further quantified as mean ± standard error in figure legends. Statistical tests are noted throughout figure legends. Results were considered significant at alpha value of p < 0.05. For multiple comparisons tests used to compare several conditions, no indication of significance in the figure means not significant. Statistical analysis was performed using Prism 9 (Graphpad software). For electrophysiology and imaging experiments, acute slices were typically generated blind to genotype and experiments were interleaved between two or three genotypes/injection conditions to control for recording conditions. Group sample sizes were chosen based on standards in the field and previous similar experiments conducted by our group. No statistical methods were used to predetermine sample size.

## ACKNOWLEDGEMENTS

We are grateful to the Bender and Ahituv lab members for comments and discussions on this manuscript. This work was supporting by SFARI grants 629287 (KJB, NA) and 513133 (KJB); the Broad Institute Target Practice Initiative (KJB); the Autism Science Foundation (ST); the Weill Neurohub Investigator Program (KJB, NA); an NSERC Predoctoral Fellowship (PWES); a Ford Foundation Dissertation Fellowship (SSH); a Weill Foundation Graduate Student Fellowship (SSH); and the following grants from the National Institute of Health: R01 MH125978 (KJB), F32 MH125536 (ADN), R01 NS078118 (JTP), R01 NS121287 (JTP), R01 MH115045 (JQP), R01 NS108874 (JQP), R01 MH118298 (JQP), T32 GM007449 (SSH).

## AUTHOR CONTRIBUTIONS

Conceptualization—KJB, SJS, NA; Methodology—ST, ADN, PWES, HK, JZ, NM, JZP, JTP, RBS, NA, KJB; Software—HK, JZ, RBS; Formal analysis—ST, ADN, PWES, HK, JZ, KJB; Investigation—ST, ADN, PWES, HK, XZ, JZ, SSH, AS, CMK, CL, ECH, KJB; Writing - Original Draft—ST, ADN; Writing - Review & Editing—All authors; Visualization—ST, ADN, PWES, HJ, JZ, KJB; Supervision—NA, KJB; Funding acquisition—ST, ADN, PWES, SSH, JTP, JZP, SJS, NA, KJB

## COMPTETING INTERESTS

NM is the cofounder, board member and CSO of Regel Therapeutics Inc and NA is the cofounder and on the scientific advisory board, Regel Tx. PWES is a scientist at Regel Tx. NM and NA are the inventors on patent ‘Gene therapy for haploinsufficiency’ WO2018148256A9. NA, KJB, and SJS receive funding from BioMarin Pharmaceutical Incorporated.

## DATA AND MATERIAL AVAILABILITY

All RNA-seq data is available on the NCBI Gene Expression Omnibus as Bioproject GSE193605. All data and analysis code are available upon request. Requests for materials can be directed to NA (CRT-based approaches) or KJB (mouse models).

## SUPPLEMENTARY MATERIALS

Figures S1 to S7

Tables S1 to S3

