## Supplemental Materials for "CRISPR activation rescues abnormalities in *SCN2A* haploinsufficiency-associated autism spectrum disorder"

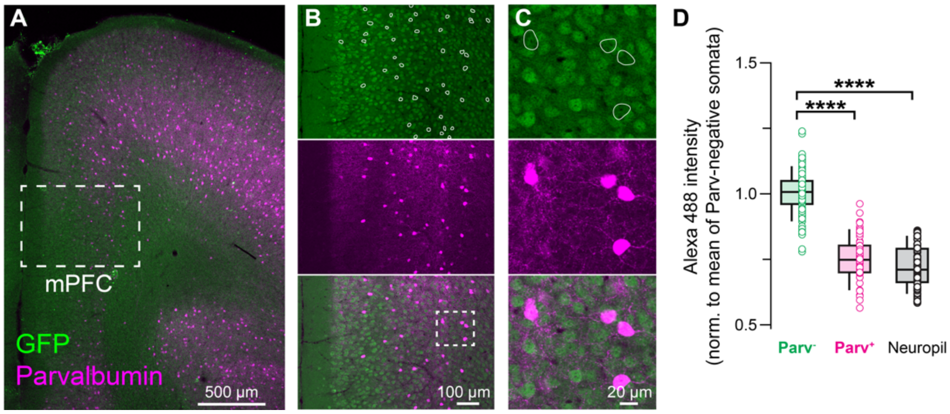

**Figure S1: Excitatory pyramidal neurons in the mPFC are GFP+ in Cre-negative *Scn2a*$^{+/KI}$ animals.**

**A-C:** Coronal brain sections from P60 *Scn2a*$^{+/KI}$ mouse (Cre-) immunostained with anti-GFP and anti-parvalbumin (PV).

**D:** Quantification of mean fluorescence intensify of GFP in PV-negative cells, PV-positive cells, and neuropil (area without somata as a measure of background fluorescence). Data are from 2 mice. Parv-: 1.0 ± 0.01, n = 67 cells; Parv+ 0.7 ± 0.01, n = 37 cells; neuropil: 0.75 ± 0.01. Parv- vs. Parv+: ****p < 0.0001. Parv- vs. neuropil: ****p < 0.0001. Holm-Šídák multiple comparisons test.

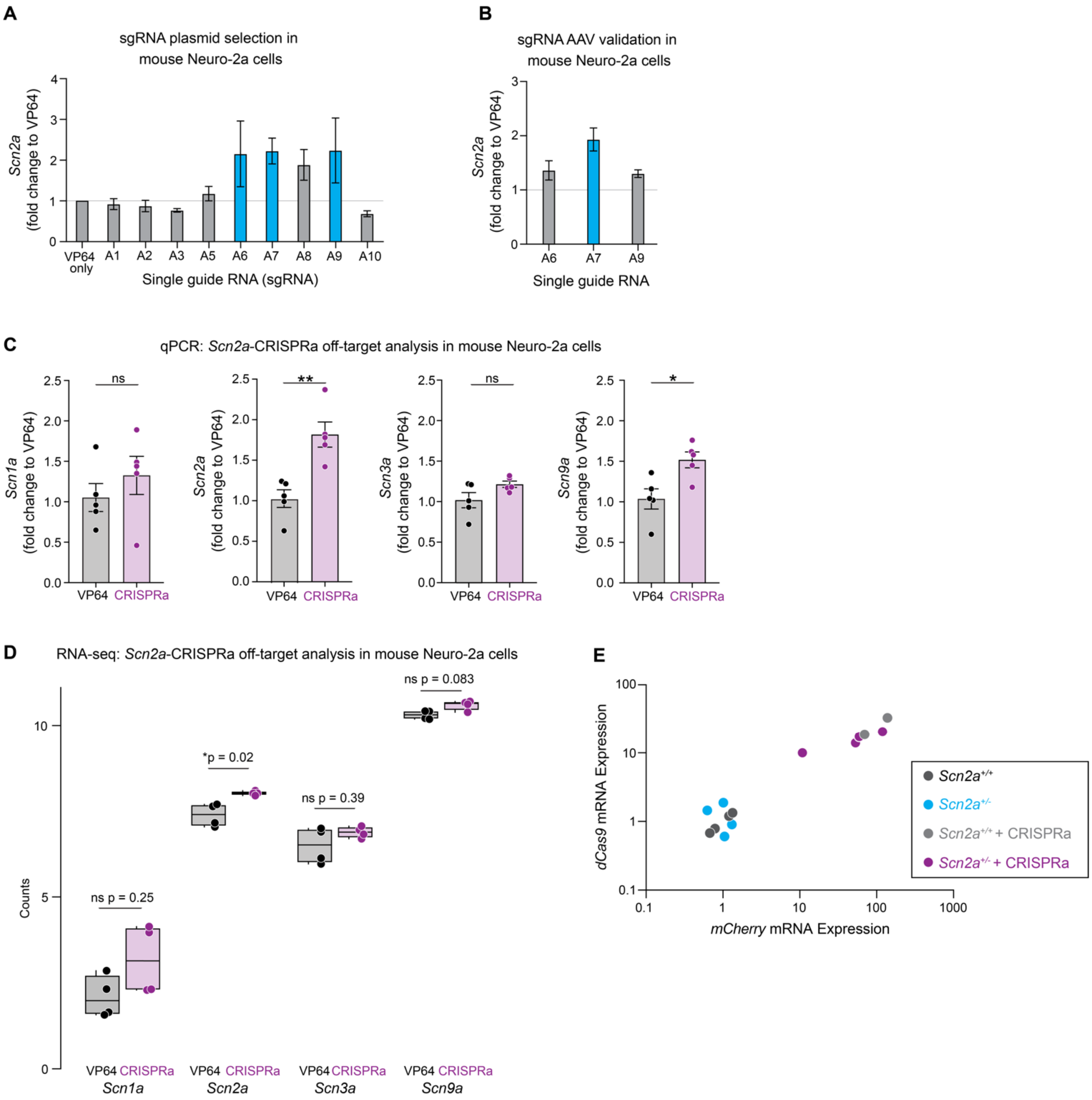

**Figure S2:** ***In vitro*** **optimization of CRISPRa constructs in mouse Neuroblastoma-2A (Neuro-2a) cells**

**A:** Fold change of *Scn2a* expression in Neuro-2a cells transfected with plasmids containing sgRNAs targeting the promoter of mouse *Scn2a* compared to a no-sgRNA VP64 control. Blue bars represent plasmids with largest increase in *Scn2a* expression.

**B:** Fold change of *Scn2a* transduced with rAAV-DJ virus in Neuro-2a cells.

**C:** qPCR off-target analysis of transcripts encoding other sodium channels in the topologically associated domain (TAD). Mann-Whitney test.

**D:** RNA-seq of other sodium channel subtypes from *Scn2a*-rAAV-CRISPRA treated Neuro-2a cells compared to VP64-only. Significance noted above data. Wald-log test.

**E:** qPCR analysis of *dCas9* and *mCherry* mRNA within the mPFC of tail vein injected *Scn2a*$^{+/+}$ + CRISPRa (light gray) and *Scn2a*$^{+/-}$ + CRISPRa (purple) versus uninjected controls *Scn2a*$^{+/+}$ (dark gray) or *Scn2a*$^{+/-}$ (cyan). Injected animals with at least a 10-fold increase in expression levels of both *dCas9* and *mCherry* to the average *Scn2a*$^{+/+}$ uninjected controls were included in EEG datasets in Fig 3.

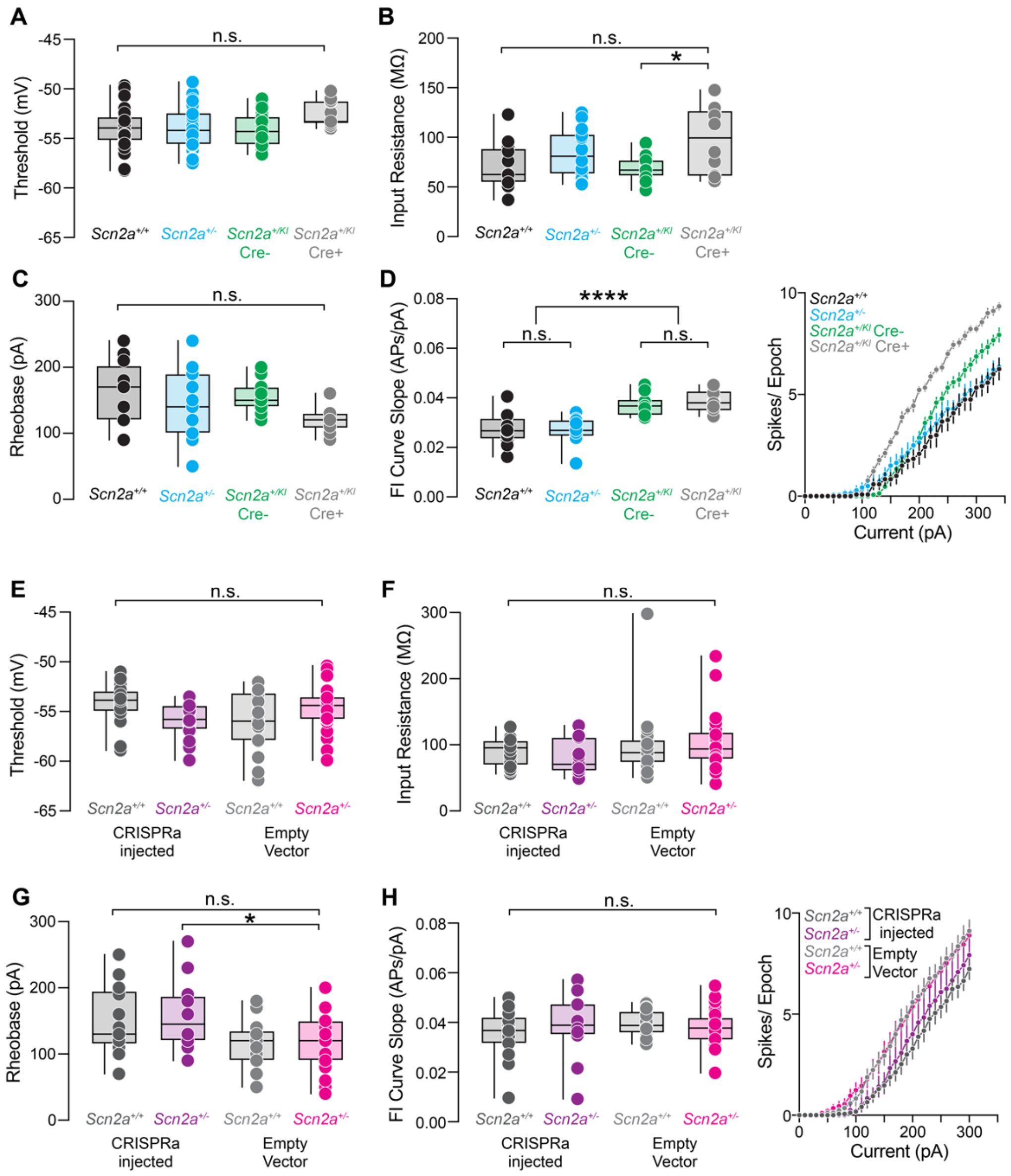

**Figure S3: Additional intrinsic electrophysiological measurements in the *Scn2a*$^{+/KI}$ conditional mouse model and CRISPRa treated neurons.**

**A:** Summary of AP threshold from P60-70 *Scn2a*$^{+/+}$ (black), *Scn2a*$^{+/-}$ (cyan), *Scn2a*$^{+/KI}$ Cre- (green) and *Scn2a*$^{+/KI}$ Cre+ (gray) neurons. Threshold of the first AP evoked by a near-rheobase current. Box plots are median and quartiles with min. and max. tails. Circles represent single cells. *Scn2a*$^{+/+}$: -54.0 ± 0.3, n = 44 cells; *Scn2a*$^{+/-}$: -54.0 ± 0.4, n = 37 cells, *Scn2a*$^{+/KI}$ Cre-: -54.1 ± 0.4 n = 18 cells; *Scn2a*$^{+/KI}$ Cre+: -52.5 ± 0.4, n = 11 cells. No significant differences. Holm-Šídák multiple comparisons test.

**B:** Summary of input resistance (MΩ). *Scn2a*$^{+/+}$: 71.3 ± 6.9, n = 12 cells; *Scn2a*$^{+/-}$: 83.8 ± 6.5, n = 14 cells, *Scn2a*$^{+/KI}$ Cre-: -69.6 ± 3.1 n = 15 cells; *Scn2a*$^{+/KI}$ Cre+: 97.7 ± 10.7, n = 10 cells. No significant differences. Holm-Šídák multiple comparisons test.

**C:** Summary of rheobase current (pA) to generate first spike. *Scn2a*$^{+/+}$: 163.3 ± 13.3, n = 12 cells; *Scn2a*$^{+/-}$: 144.3 ± 13.6, n = 14 cells, *Scn2a*$^{+/KI}$ Cre-: 153.3 ± 5.6 n = 15 cells; *Scn2a*$^{+/KI}$ Cre+: 119.0 ± 6.1, n = 10 cells. No significant differences. Holm-Šídák multiple comparisons test.

**D:** APs per 300 ms stimulation epoch for each current amplitude. Left: Quantification of firing rate slope of data on Right. *Scn2a*$^{+/+}$: 0.03 ± 0.002, n = 12 cells; *Scn2a*$^{+/-}$: 0.03 ± 0.002, n = 14 cells, *Scn2a*$^{+/KI}$ Cre-: 0.04 ± 0.001, n = 15 cells; *Scn2a*$^{+/KI}$ Cre+: 0.04 ± 0.001, n = 10 cells. *Scn2a*$^{+/+}$ vs. *Scn2a*$^{+/KI}$ Cre-: ****p < 0.0001, *Scn2a*$^{+/+}$ vs. *Scn2a*$^{+/KI}$ Cre+: ****p < 0.0001, *Scn2a*$^{+/-}$ vs. *Scn2a*$^{+/KI}$ Cre-: ****p < 0.0001. *Scn2a*$^{+/-}$ vs. *Scn2a*$^{+/KI}$ Cre+: ****p < 0.0001. Holm-Šídák multiple comparisons test. Right: Number of APs versus current amplitude injected.

**E:** Summary of AP threshold from P57-85 *Scn2a*-rAAV-CRISPRa treated *Scn2a*$^{+/+}$ (dark gray) and *Scn2a*$^{+/-}$ (purple) neurons and *Scn2a*-rAAV-empty transduced *Scn2a*$^{+/+}$ (light gray) and *Scn2a*$^{+/-}$ (magenta) neurons. Threshold of the first AP evoked by a near-rheobase current. represent single cells. *Scn2a*$^{+/+}$ + CRISPRa: -54.2 ± 0.4, n = 24 cells; *Scn2a*$^{+/-}$

+ CRISPRa: -55.7 ± 0.4, n = 19 cells, *Scn2a*$^{+/+}$ + empty: -56 ± 0.7 n = 18 cells; *Scn2a*$^{+/-}$ + empty: -55.6 ± 0.4, n = 29 cells. No significant differences. Holm-Šídák multiple comparisons test.

**F:** Summary of input resistance (MΩ) from *Scn2a*-rAAV-CRISPRa or *Scn2a*-rAAV-empty neurons. *Scn2a*$^{+/+}$ + CRISPRa: 89.5 ± 5.0, n = 17 cells; *Scn2a*$^{+/-}$ + CRISPRa: 80.5 ± 8.0, n = 11 cells, *Scn2a*$^{+/+}$ + empty: 99.3 ± 12.7, n = 18 cells; *Scn2a*$^{+/-}$ + empty: 97.7 ± 10.7, n = 29 cells. No significant differences. Holm-Šídák multiple comparisons test.

**G:** Summary of rheobase current (pA) to generate first spike from *Scn2a*-rAAV-CRISPRa or *Scn2a*-rAAV-empty neurons. *Scn2a*$^{+/+}$ + CRISPRa: 148.2 ± 11.6, n = 17 cells; *Scn2a*$^{+/-}$ + CRISPRa: 157.5 ± 15.2, n = 12 cells, *Scn2a*$^{+/+}$ + empty: 116.5 ± 8.5, n = 17 cells; *Scn2a*$^{+/-}$ + empty: 115.9 ± 7.8, n = 29 cells. *Scn2a*$^{+/-}$ + CRISPRa vs. *Scn2a*$^{+/-}$ + empty: *p = 0.04. Holm-Šídák multiple comparisons test.

**H:** APs per 300 ms stimulation epoch for each current amplitude. Left: Quantification of firing rate slope of data on Right. *Scn2a*$^{+/+}$ + CRISPRa: 0.04 ± 0.002, n = 17 cells; *Scn2a*$^{+/-}$ + CRISPRa: 0.04 ± 0.004, n = 12 cells, *Scn2a*$^{+/+}$ + empty: 0.04 ± 0.001, n = 17 cells; *Scn2a*$^{+/-}$ + empty: 0.04 ± 0.001, n = 29 cells. No significant differences. Holm-Šídák multiple comparisons test. Right: Number of APs versus current amplitude injected.

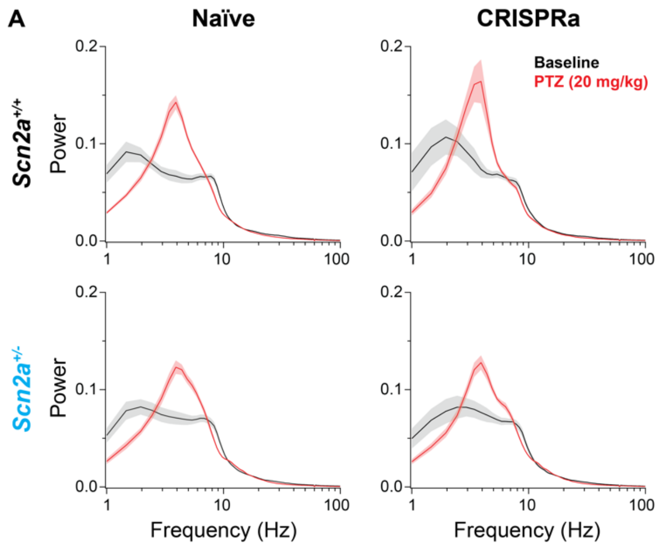

**Figure S4: Power spectral density (PSD) during baseline and 20 mg/kg PTZ administration**

**A:** Average PSD across *Scn2a*$^{+/+}$: n = 24 mice; *Scn2a*$^{+/-}$: n = 16 mice; *Scn2a*$^{+/+}$ + CRISPR: n = 7 mice; *Scn2a*$^{+/-}$ + CRISPRa: n = 12 mice during baseline (black) and 20 mg/kg PTZ (red). Note marked increase in PSD in 3-5 Hz observed across all conditions.

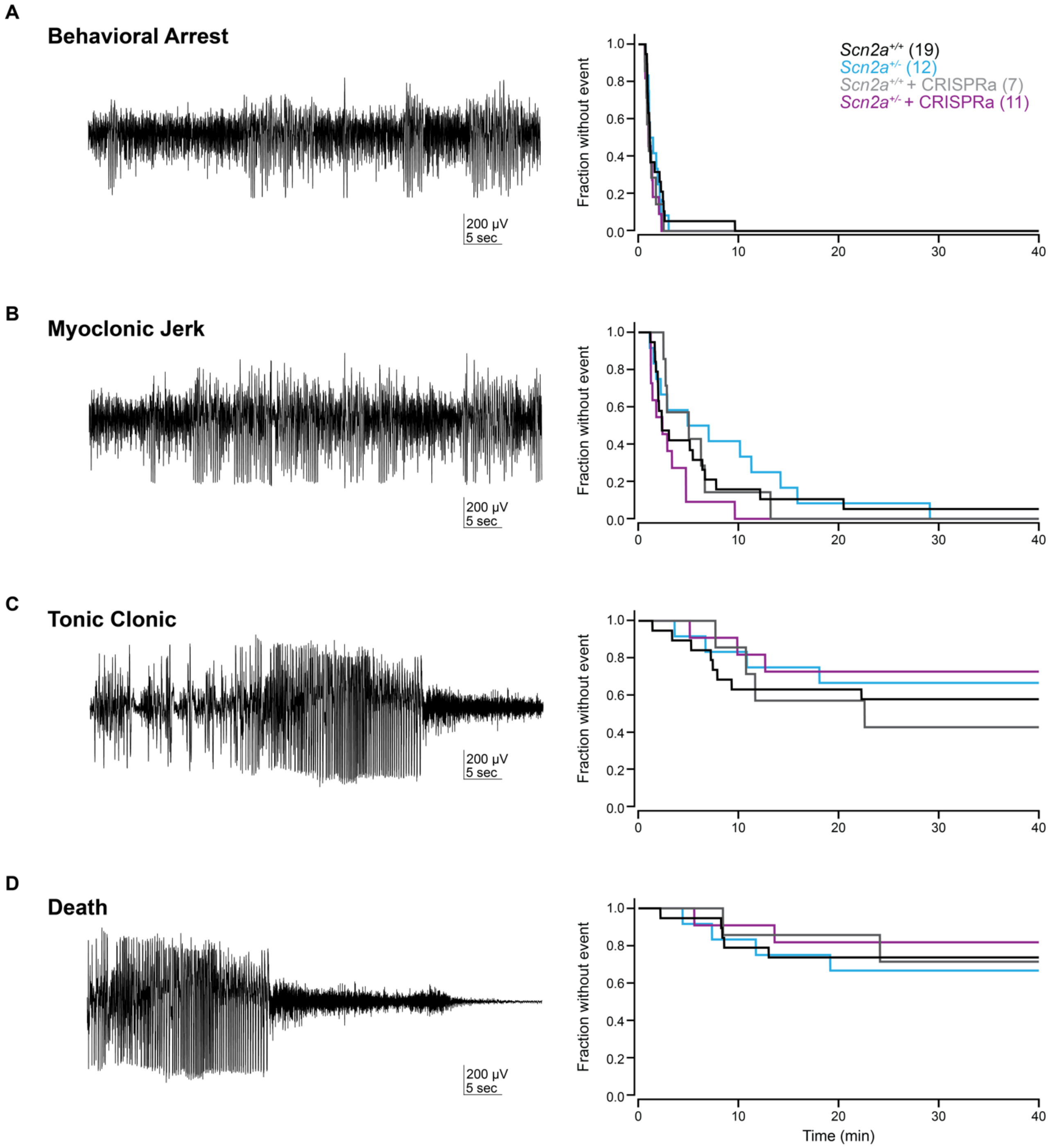

**Figure S5: Detection of 50 mg/kg PTZ-induced seizures**

**A-D:** Left: Example EEG trace of behavioral arrest ($p = 0.51$), myoclonic jerk ($p = 0.14$), tonic clonic seizure ($p = 0.69$; reprinted data from Main Fig. 3), and death ($p = 0.65$, Mantel log-rank test). Right: Survival curves over 40 minutes across *Scn2a*$^{+/+}$ (black): n = 19 mice; *Scn2a*$^{+/-}$ (cyan): n = 12 mice; *Scn2a*$^{+/+}$ + CRISPRa (gray): n = 7 mice; *Scn2a*$^{+/-}$ + CRISPRa (purple): n = 11.

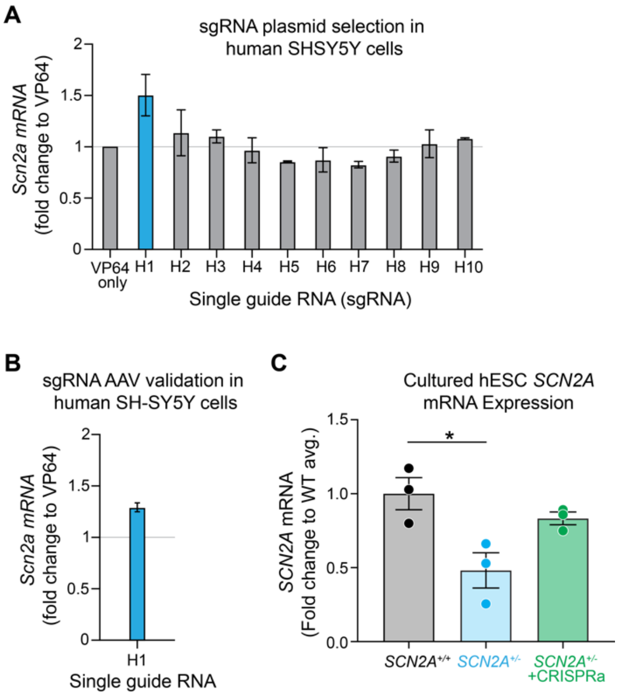

**Figure S6: *In vitro* optimization of CRISPRa constructs in human SH-SY5Ycells**

**A:** Fold change of *Scn2a* expression in SH-SY5Y cells transfected with plasmids containing sgRNAs targeting the promoter of human *Scn2a* compared to a no sgRNA VP64 control.

**B:** Fold change of *Scn2a* transduced with rAAV-DJ virus in human SH-SY5Y cells.

**C:** *SCN2A* mRNA expression from *SCN2A*$^{+/+}$ (black), *SCN2A*$^{+/-}$ (cyan), and *SCN2A*-rAAV-CRISPRa treated *SCN2A*$^{+/-}$ (purple) hESC-derived neurons normalized to wild type average. *SCN2A*$^{+/+}$: 1.0 ± 0.1, n = 3 dishes; *SCN2A*$^{+/-}$: 0.48 ± 0.1, n = 3 dishes; *SCN2A*$^{+/-}$ + CRISPRa: 0.8 ± 0.04, n = 3 dishes. *SCN2A*$^{+/+}$ vs. *SCN2A*$^{+/-}$: *p = 0.03. Holm-Šídák multiple comparisons test.

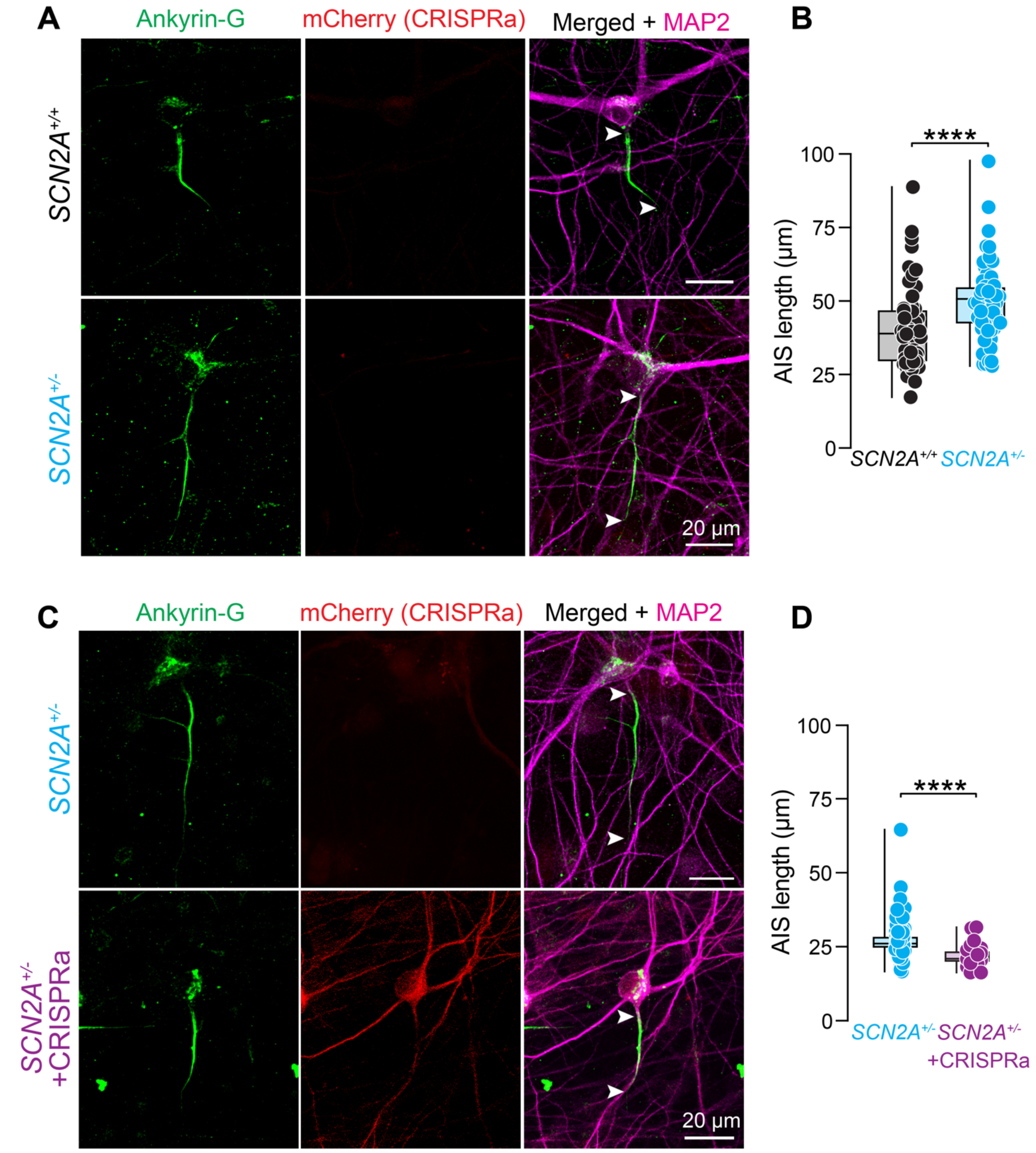

**Figure S7: Axon initial segment structural plasticity in *SCN2A*$^{+/-}$ neurons is rescued by CRISPRa.**

**A:** Representative images of *SCN2A*$^{+/+}$ (black) and *SCN2A*$^{+/-}$ (cyan) human stem-cell-derived neurons immunostained with antibodies against ankyrin-G (green) and MAP2 (magenta). Arrows denote start and end points used to quantify AIS length.

**B:** Quantification of AIS length. *SCN2A*$^{+/+}$: 40.5 ± 1.9 µm, n = 56 cells, 3 dishes. *SCN2A*$^{+/-}$: 50.2 ± 1.8 µm, n = 56 cells, 3 dishes. ****p < 0.0001. Mann-Whitney test.

**C:** Representative images of *SCN2A*$^{+/-}$ neurons expressing *Scn2a*-rAAV-CRISPRa-mCherry (purple) and mCherry-negative internal *SCN2A*$^{+/-}$ controls (cyan). Immunostaining against ankyrin-G and MAP2.

**D:** Quantification of AIS length. *SCN2A*$^{+/-}$: 27.1 ± 0.5 µm, n = 122 cells, 3 dishes. *SCN2A*$^{+/-}$ + CRISPRa: 22.0 ± 0.8 µm, n = 23 cells, 3 dishes. ****p < 0.0001. Mann-Whitney test.

**Supplemental Table 1 (Attached): ASD risk genes (Fu et al., 2021) and haploinsufficiency likelihood.**
All genes shown with gene IDs, cDNA length, presence in ASD72 list (genes with genome-wide significance, Fu et al., 2021), and pLI (probability of being loss-of-function intolerance) and LOEUF (loss-of-function observed/expected upper bound fraction) scores. Genes in blue have cDNA > 3000 BPs with a LOEUF score of < 0.273.

**Supplemental Table 2: sgRNA sequences tested per species, and primer sequences used for qPCR.**

| **Primers** | **Forward** | **Reverse** |
|---|---|---|
| qPCR-Mouse_Scn2a | ATTTTCGGCTCATTCTTCACACT | GGGCGAGGTATCGGTTTTTGT |
| qPCR-Mouse_Bactin | GACGATGCTCCCCGGGCTGTATTC | TCTCTTGCTCTGGGCCTCGTCACC |
| qPCR-sadCas9-VP64 | ATCACCCCCCACCAGATCAAGC | GTCCTTGTCGTACAGGCCGTTCA |
| qPCR-Human_*SCN2A* | CGCTTCTTTACCAGGGAATCC | TCCTGTTTGGGTCTCTTAGCTTT |
| qPCR-Human_Bactin | CATGTACGTTGCTATCCAGGC | CTCCTTAATGTCACGCACGAT |
| genotyping-Mouse_Scn2a | TGCGAGGAGCTAAACAGTGATTAAAG | GGCTCCATTCCCTTATCAGACCTACCC |
| **tested sgRNA** | | |
| mouse_*Scn2a* | sgRNA sequence | |
| A1 | ACAGAATCAGTAACGCACTGT | |
| A2 | CGGGTAAGCCAAGTTTAGTCA | |
| A3 | AAGCACTTGCCTCACATAAAT | |
| A5 | CTAGGTCATAGAAAGGAAACC | |
| A6 | TTTATTGGACCCCAGATATTC | |
| A7 | AGAAAATTAACTTAGTGCATA | |
| A8 | AAGCCGCCAGGGACCCGAGCA | |
| A9 | TATAACTGCCACTAGAGGGCT | |
| A10 | GACCCTCCTCCGGGCTCCACC | |
| human_*SCN2A* | sgRNA sequence | |
| H1 | TGCTGACTGCTACATAGCCAA | |
| H2 | GTGCTGACTGCTACATAGCCA | |
| H3 | CTGCTACATAGCCAAAGGAAC | |
| H4 | GCTCCATCTCCTGGTCAAAAG | |
| H5 | CAGCCCATAATTCCACTCTAT | |
| H6 | AGTAGTTGATTTCAAATAGAG | |
| H7 | ATTAAAGTAGTTGATTTCAAA | |
| H8 | GATTTCAAATAGAGTGGAATT | |
| H9 | AAGTAGTTGATTTCAAATAGA | |
| H10 | AGCTCCATCTCCTGGTCAAAA | |

**Supplemental Table 3: AAV titers for all viruses used**

| **Sequence** | **Serotype** | **Plasmid** | **Genomic Titer (vg/mL)** |
|---|---|---|---|
| Mouse sgRNA | DJ | pU6-sasgRNA-CMV-mCherry | 4.40E+13 |
| Human sgRNA | DJ | pU6-sasgRNA-CMV-mCherry | 1.33E+14 |
| mCherry | DJ | pAAV-CMV-mCherry | 3.20E+13 |
| sadCas9VP64 | DJ | pCMV-sadCas9-VP64-pA | 4.82E+13 |
| Mouse sgRNA | PhP.eb | pU6-sasgRNA-CMV-mCherry | 5.01E+13 |
| mCherry | PhP.eb | pAAV-CMV-mCherry | 5.00E+13 |
| sadCas9VP64 | PhP.eb | pCMV-sadCas9-VP64-pA | 5.02E+13 |
